# A druggable oxidative folding pathway in the endoplasmic reticulum of human malaria parasites

**DOI:** 10.1101/2020.05.13.093591

**Authors:** David W. Cobb, Heather M. Kudyba, Alejandra Villegas, Michael R. Hoopmann, Rodrigo Baptista, Baylee Bruton, Michelle Krakowiak, Robert L. Moritz, Vasant Muralidharan

## Abstract

Malaria remains a major global health problem, and there exists a constant need to identify druggable weaknesses in *P. falciparum* biology. The endoplasmic reticulum (ER) has many essential roles in the asexual lifecycle and may offer new drug targets, but it remains critically understudied. We generated conditional mutants of the putative redox-active, ER chaperone *Pf*J2, and show that it is essential for parasite survival. Using a redox-active cysteine crosslinker, we identify its substrates to be other mediators of oxidative folding, *Pf*PDI8 and *Pf*PDI11, suggesting a redox-regulatory role for *Pf*J2. Knockdown of these protein disulfide isomerases in *Pf*J2 conditional mutants show that *Pf*PDI11 is not essential, while *Pf*PDI8 is essential for asexual growth and may work in a complex with PfJ2 and other ER chaperones. Finally, we show that these redox interactions in the parasite ER are sensitive to small molecule inhibition. Together these data build a model for how oxidative folding occurs in the *P. falciparum* ER and demonstrate its suitability for antimalarial drug development.

## Introduction

Today, the majority of the world’s population lives at risk for contracting malaria, a disease caused by eukaryotic parasites of the genus *Plasmodium*, with *P. falciparum* causing the most severe forms of the disease (World Heath Organization, 2019). In 2018, the world saw approximately 228 million cases of malaria resulting in more than 400,000 deaths. These numbers reflect a concerted effort to combat malaria in the past few decades, but progress has stagnated, with the numbers of malaria cases/deaths largely unchanged in recent years. A major impediment in the fight against malaria is the rise of drug-resistant— including multidrug-resistant—*P. falciparum* parasites, highlighting the need for more research into the biology of this major human pathogen.

The morbidity and mortality of malaria is a direct result of asexual parasite replication inside of human red blood cells (RBCs) (Cowman et al., 2016). As such, many validated and proposed drug targets involve the processes that support this lifecycle: these include parasite signaling events, egress/invasion, metabolism, plastid function, and remodeling of the RBC (Absalon et al., 2018; Amberg-Johnson et al., 2017; Bowman et al., 2014; Favuzza et al., 2020; Hodder et al., 2015; Moura et al., 2009; Nasamu et al., 2017; Pino et al., 2017). Strikingly, all of these diverse aspects of *P. falciparum* biology, each promising drug targets on their own, are united in their reliance on the parasite secretory pathway that originates in the endoplasmic reticulum (ER). Therefore, the *P. falciparum* ER is likely an Achilles’ heel, with inhibition of its function having far-reaching effects on parasite biology. In fact, the frontline antimalarial artemisinin and its derivatives were shown in recent years to induce ER stress, indicating that the *P. falciparum* ER is sensitive to drug treatment (Bridgford et al., 2018; Zhang et al., 2017). Additionally, a promising new class of antimalarial drugs has been shown to target ER function (LaMonte et al., 2020). And yet, despite being a potential drug target in its own right, the parasite ER remains critically understudied.

Proteins are co-translationally imported into the ER and must correctly fold, or risk inducing ER stress and cell death (Almanza et al., 2019). One aspect of this process is the “oxidative folding” that occurs within the ER—the formation and reduction of disulfide bonds in proteins as they try to achieve their native states in the ER’s oxidizing environment. ER-resident, Thioredoxin (Trx) superfamily proteins catalyze oxidative folding and are therefore essential for ER function (Hatahet & Ruddock, 2009). Highlighting their importance, these ER Trx-domain proteins are proposed drug targets in other human diseases (Hoffstrom et al., 2010; Kaplan et al., 2015; Vatolin et al., 2016; Xu et al., 2012). How oxidative folding and Trx-domain proteins work in the *P. falciparum* ER is unknown, and the parasite encodes far fewer ER-resident members of the Trx superfamily relative to model eukaryotic systems (Galligan & Petersen, 2012; Mahajan et al., 2006; Mouray et al., 2007). This streamlined approach to oxidative folding suggests less redundancy between the family members: elucidation of their functions will reveal new insights into parasite biology and potentially provide novel targets for antimalarial drug development.

We report here that a putative ER Trx-domain-containing chaperone *Pf*J2 is essential for the *P. falciparum* asexual lifecycle. We identify PfJ2 interacting partners and use a chemical biology approach to specifically identify its redox partnerships. We show that one such partner—*Pf*PDI8, also a Trx-superfamily member—is another essential ER protein that mediates oxidative folding, and our data suggests that both proteins work together with other chaperones to promote protein folding in the ER. Finally, we demonstrate that the redox interactions between these essential proteins and their substrates are amenable to disruption by a small molecule inhibitor, and we suggest that the oxidative folding process of the *P. falciparum* ER is an exploitable drug target.

## Results

### *Pf*J2 is an essential, ER-resident Hsp40

*Pf*J2 is a putative ER-resident Hsp40 with a C-terminal thioredoxin (Trx) domain (Figure 1A, B). It has similarity to a protein in the mammalian ER which has several Trx domains following an Hsp40 J-domain, suggesting PfJ2 may catalyze disulfide bond reduction of client proteins (Cunnea et al., 2003; Oka et al., 2013; Ushioda et al., 2008) (Figure 1A). To investigate *Pf*J2 function and its potential role in *P. falciparum* oxidative folding, we generated a *Pf*J2 conditional knockdown parasite line using the TetR-*Pf*DOZI aptamer system (referred to as *Pf*J2_apt_ hereafter). In this knockdown system, protein expression is regulated by the presence of anhydrotetracycline (aTc), with knockdown induced by removal of aTc (Ganesan et al., 2016) (Figure 1C). Using CRISPR/Cas9 genome editing, we modified the *pfj2* locus to encode a 3xHA-tag immediately upstream of the ER-retention signal, as well as the regulatory aptamer sequences and a cassette to express the TetR-*Pf*DOZI fusion protein (Figure 1D). Correct integration of the construct into the *pfj2* locus was determined by PCR, and expression of HA-tagged *Pf*J2 was demonstrated via western blot (Figure 1D).

**Figure 1.**
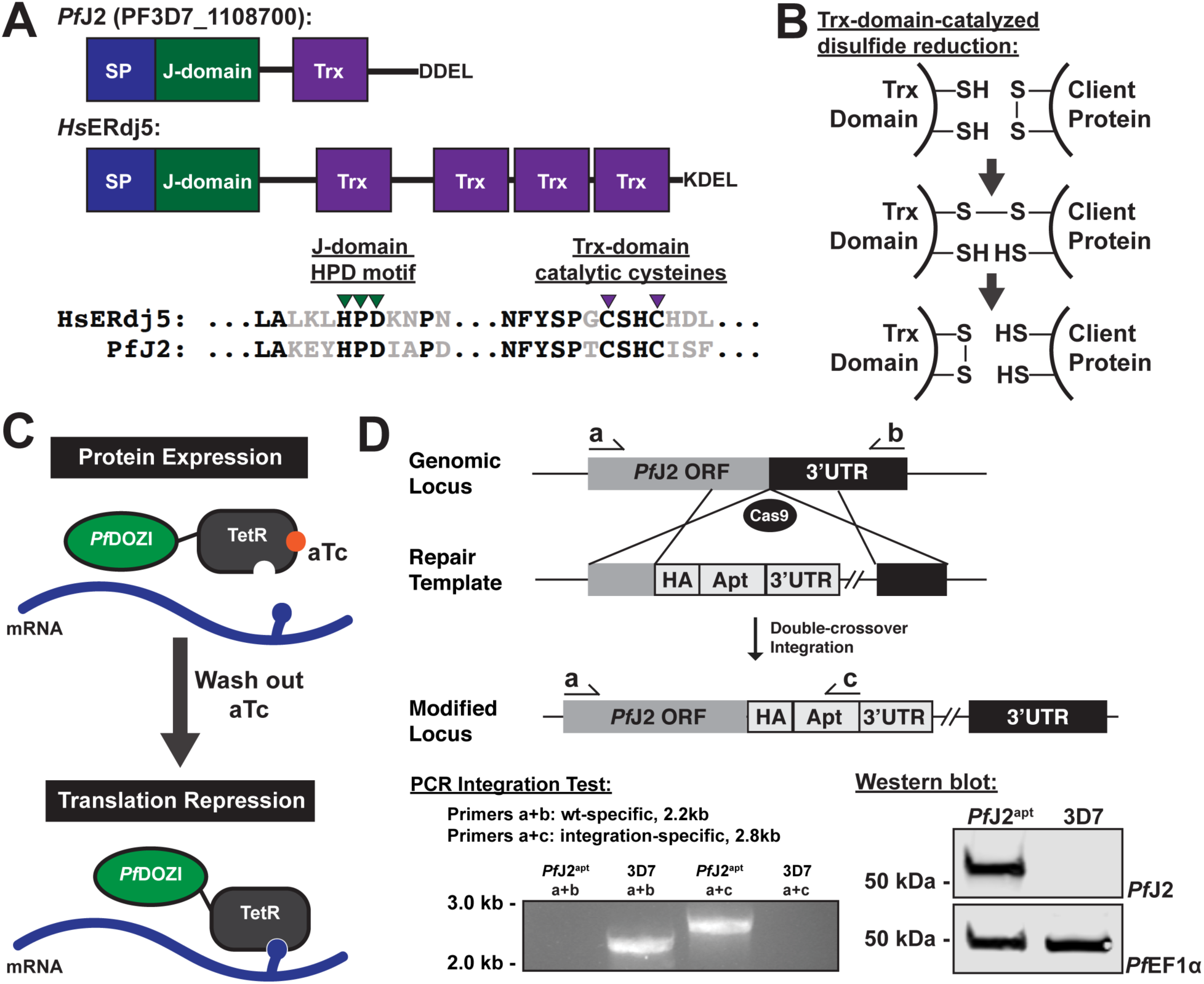
Generation of *Pf*J2 (PF3D7_1108700) conditional knockdown mutants using CRISPR/Cas9. **A)** Predicted domain structure of *Pf*J2 and human homolog ERdj5, showing signal peptide (SP), Hsp40 J-domain, thioredoxin domain (Trx), and C-terminal ER retention signals. Essential, conserved residues are shown: the J-domain HPD motif is required for Hsp40 activity (i.e., stimulation of Hsp70 ATPase activity), and the Trx-domain CXXC motif is required for redox activity. Shown is homology between the PfJ2 Trx-domain and the first ERdj5 Trx-domain active sites. **B)** Mechanism of disulfide bond reduction catalyzed by Trx-domain active site cysteines. **C)** Regulation of protein expression using the TetR-*Pf*DOZI knockdown system. TetR binds to aptamer sequences present in the mRNA, and *Pf*DOZI localizes the complex to sites of mRNA sequestration, repressing translation. Anhydrotetracycline (aTc) blocks TetR-aptamer interaction. **D)** Schematic of CRISPR/Cas9-mediated introduction of the TetR-*Pf*DOZI knockdown system into the *pfj2* locus. A linearized repair template was transfected, along with a plasmid to express Cas9 and a gRNA, to introduce sequences for a 3xHA tag+*Pf*J2’s DDEL ER retention signal, stop codon, and a 3’UTR. Included in the repair template but not shown was a cassette to express the TetR-*Pf*DOZI fusion protein and blasticidin deaminase for drug selection. Bottom left: two PCR integration tests were used to amplify either a sequence from only wild-type locus (primers a+b) or the modified locus (a+c). Bottom right: anti-HA western blot.

Using an immunofluorescence assay (IFA), we assessed the localization of *Pf*J2 throughout the asexual lifecycle (Figure 2A). We were especially keen to determine PfJ2’s localization given the presence of an apparent Plasmodium Export Element (PEXEL motif) downstream of the signal peptide (Botha et al., 2007). A processed PEXEL motif would mark PfJ2 for export into the host RBC, in direct contrast to the “XXEL” ER retention signal at the C-terminus (Hiller, 2004; Külzer et al., 2009; Marti, 2004; Ilaria Russo et al., 2010). Co-localization between *Pf*J2 and the ER-marker Plasmepsin V (*Pf*PMV) revealed that *Pf*J2 is in fact an ER-resident protein, and we were unable to detect *Pf*J2 signal outside of the ER by IFA (Figure 2A). Using highly synchronized parasites, we showed that *Pf*J2 is primarily expressed in the trophozoite and schizont stages, and that during knockdown conditions (removal of aTc), *Pf*J2 expression is reduced (Figure 2B). Importantly, knockdown of *Pf*J2 was found to inhibit expansion of parasites in culture, with an aTc EC_50_ of approximately 0.5 nM (Figure 2C). Consistent with peak *Pf*J2 expression during the trophozoite/schizont stages, we observed normal development of knockdown parasites in the beginning of the asexual life cycle, but the development of these parasites began to slow in the trophozoite stage and they failed to complete schizogony and produce new daughter parasites (Supplementary Figure 1, Figure 2D). These data demonstrate that *Pf*J2 is a *bona fide* ER-resident protein essential for progression through the *P. falciparum* asexual lifecycle.

**Figure 2.**
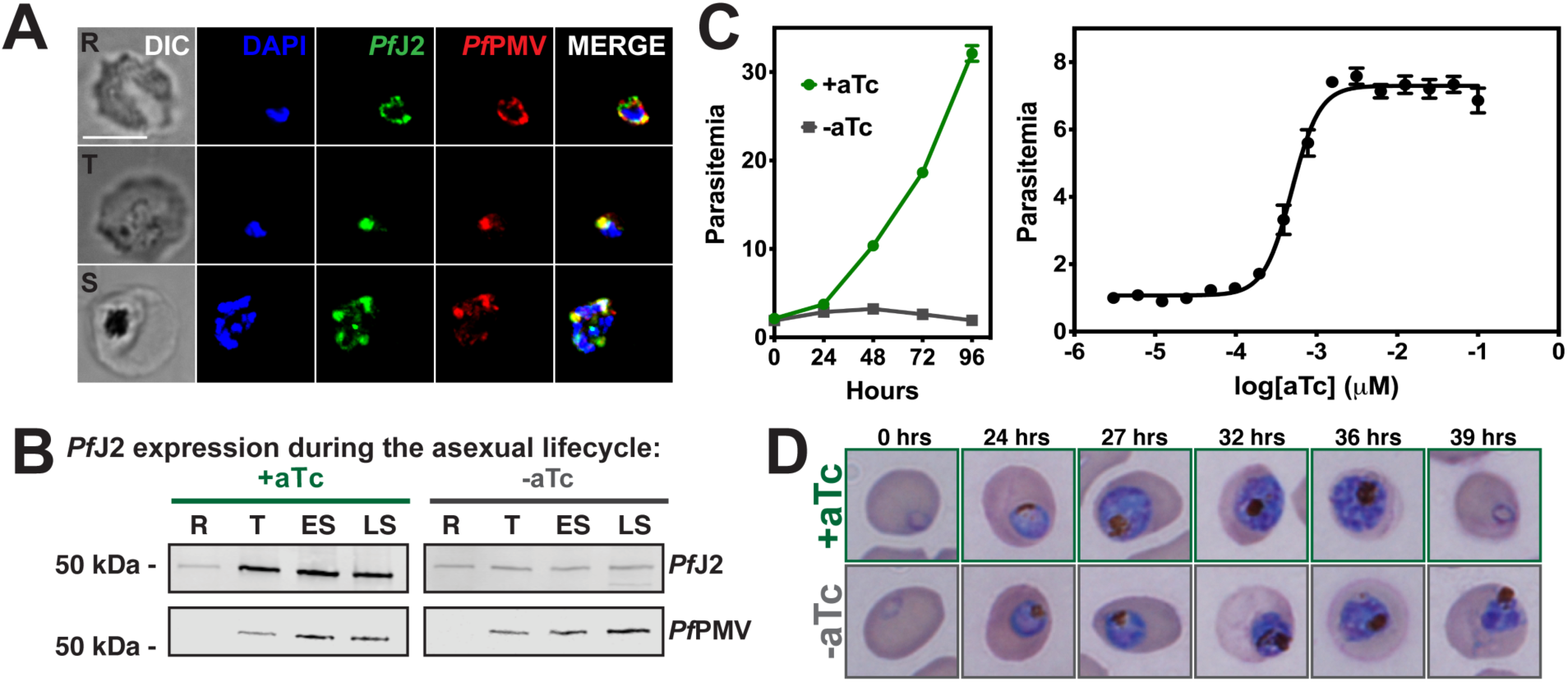
*Pf*J2 is an essential, ER-resident protein. **A)** *Pf*J2^apt^ parasites were fixed with paraformaldehyde and glutaraldehyde and stained with DAPI (blue) and with antibodies against HA (green) and the ER-marker *Pf*PMV (red). Ring (R), Trophozoite (T), and Schizont (S) stage parasites are shown. Z-stack Images were deconvoluted and shown as a single, maximum intensity projection. Scale bar represents 5 μm. **B)** Parasites were tightly synchronized to the ring stage (0-3 hours) and split into two conditions: +aTc and –aTc. Samples were taken for western blot analysis at various time points in the life cycle (R = Ring, T = Trophozoite, ES = Early Schizont, LS = Late Schizont). Equal parasite equivalents were loaded into each lane, and blots were stained with antibodies for HA and *Pf*PMV. **C)** Left: asynchronous parasites were grown in normal (+aTc) or *Pf*J2 knockdown (-aTc) conditions, and parasite growth was monitored daily for 96 hours via flow cytometry. Right: asynchronous parasites were grown in a range of aTc concentrations and growth measured at 72 hours via flow cytometry. The aTc EC_50_ was determined to be 0.5 nM. **D)** Parasites were tightly synchronized to the ring stage (0-3 hours) and split into two conditions: normal (+aTc) and PfJ2 knockdown (-aTc). Samples from each condition were smeared and field-stained at time points throughout the lifecycle.

### *Pf*J2 interacts with essential ER chaperones and proteins in the secretory pathway

As a putative chaperone possibly involved in ER oxidative folding, we reasoned that the essentiality of *Pf*J2 is likely related to its ability to interact with other proteins in the ER. We therefore took a co-immunoprecipitation (coIP) approach to identify *Pf*J2 interacting partners. *Pf*J2 was immunoprecipitated from *Pf*J2^apt^ parasite lysates using anti-HA antibodies, and co-immunoprecipitating proteins were identified by tandem mass spectrometry (MS/MS) analysis. Control parental parasites (lacking HA-tagged *Pf*J2) were also used for immunoprecipitation and analyzed in the same manner. Each co-IP experiment was performed in triplicate, and the abundance of each identified protein was calculated by summing the total MS1 intensities of all matched peptides for each selected protein, and normalizing by the total summed intensity of all matched peptides in the sample (Boucher et al., 2018; Florentin et al., 2019). Because *Pf*J2 is an ER-localized protein, we further narrowed our list of interacting partners to those containing a signal peptide and/or at least one transmembrane domain (i.e. proteins predicted to be within the ER or traffic through the secretory pathway). We identified a stringent list of high-confidence interacting partners as those proteins which were present in all three *Pf*J2^apt^ coIP experiments, and were at least 5-fold more abundant compared to the controls, as previously described (Florentin et al., 2019) (Supplementary Figure 2). A complete list of identified proteins is provided in Supplementary Table 1.

**Table 1.**
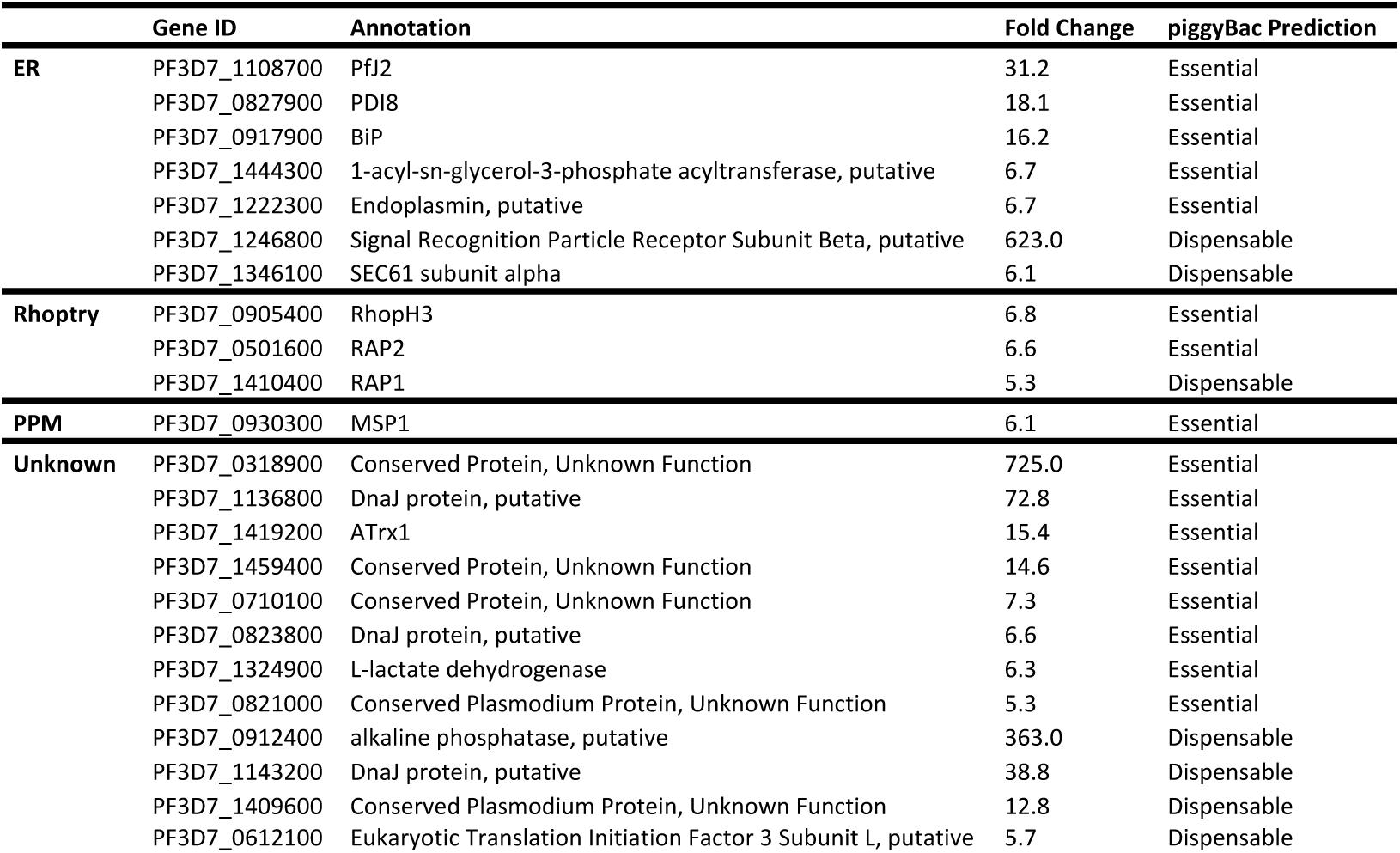
Proteins identified as PfJ2 interacting partners. Identified proteins were categorized by subcellular localization (ER, Rhoptry, Parasite Plasma Membrane [PPM], or Unknown). Also shown are GeneIDs and annotations from PlasmoDB.org, calculated fold-change compared to control experiments, and essentiality as predicted by the piggyBac mutagenesis screen performed by Zhang et al., 2018.

We identified other conserved proteins classically involved in essential ER processes— such as the Hsp70 Binding immunoglobulin Protein (BiP), the Hsp90 Endoplasmin, and the oxidoreductase Protein Disulfide Isomerase (PDI) (Table 1). We further identified proteins that are trafficked through the ER late in the parasite lifecycle and are required for egress and invasion, including PfMSP1 and proteins destined for rhoptries (Das et al., 2015; Ito et al., 2017; Richard et al., 2009; Sherling et al., 2017). Nearly half of the identified proteins lack empirical evidence for their subcellular localization, and many have no known function. But, given the presence of a signal peptide and/or transmembrane domains, these proteins likely have localizations in the ER, parasite plasma membrane, apicoplast, and other destinations that are part of the secretory pathway. Also of note, approximately two-thirds of the identified proteins are predicted to have essential functions (Supplementary Figure 2) (Zhang et al., 2018). These data together suggest that *Pf*J2 may work with other ER-resident chaperones to ensure proper folding/functioning of proteins that have essential roles throughout the parasite.

### PfJ2 is a redox-active protein in the *P. falciparum* ER

We next sought to test the hypothesis that *Pf*J2 contributes to ER oxidative folding by determining whether its putative Trx domain is redox-active and capable of forming mixed disulfides with its clients. To this end we employed the bifunctional, electrophilic crosslinker divinyl sulfone (DVSF), which has been shown in model systems to have remarkable specificity for redox-active, nucleophilic cysteines, like those present in Trx domains (Allan et al., 2016; Araki et al., 2017; Naticchia et al., 2013). This specificity allows DVSF to covalently and irreversibly trap Trx-domain proteins to their redox substrates (Figure 3A). Therefore, we treated *Pf*J2^apt^ parasite cultures with DVSF and isolated proteins for western blot analysis. In the absence of DVSF, *Pf*J2 was detected at approximately 50 kDa, while the addition of DVSF resulted in additional bands containing *Pf*J2 to appear between 100-150 kDa (Figure 3A). To demonstrate specificity of DVSF for redox-active cysteines, the ER-resident protein *Pf*PMV—which contains 16 cysteines after its signal peptide—was detected in the same samples and its migration pattern was found to be unaffected by DVSF treatment (Figure 3A). As an additional control, *Pf*J2^apt^ parasite cultures were treated with the sulfhydryl-blocking compound N-ethylmaleimide (NEM) prior to the addition of DVSF. Pre-treatment with NEM resulted in the blockage of cross-linking between *Pf*J2 and its redox partners (Figure 3B). These results demonstrate that *Pf*J2 does have redox activity, and that DVSF is a useful chemical tool for trapping redox interactions in the ER of *P. falciparum*.

**Figure 3.**
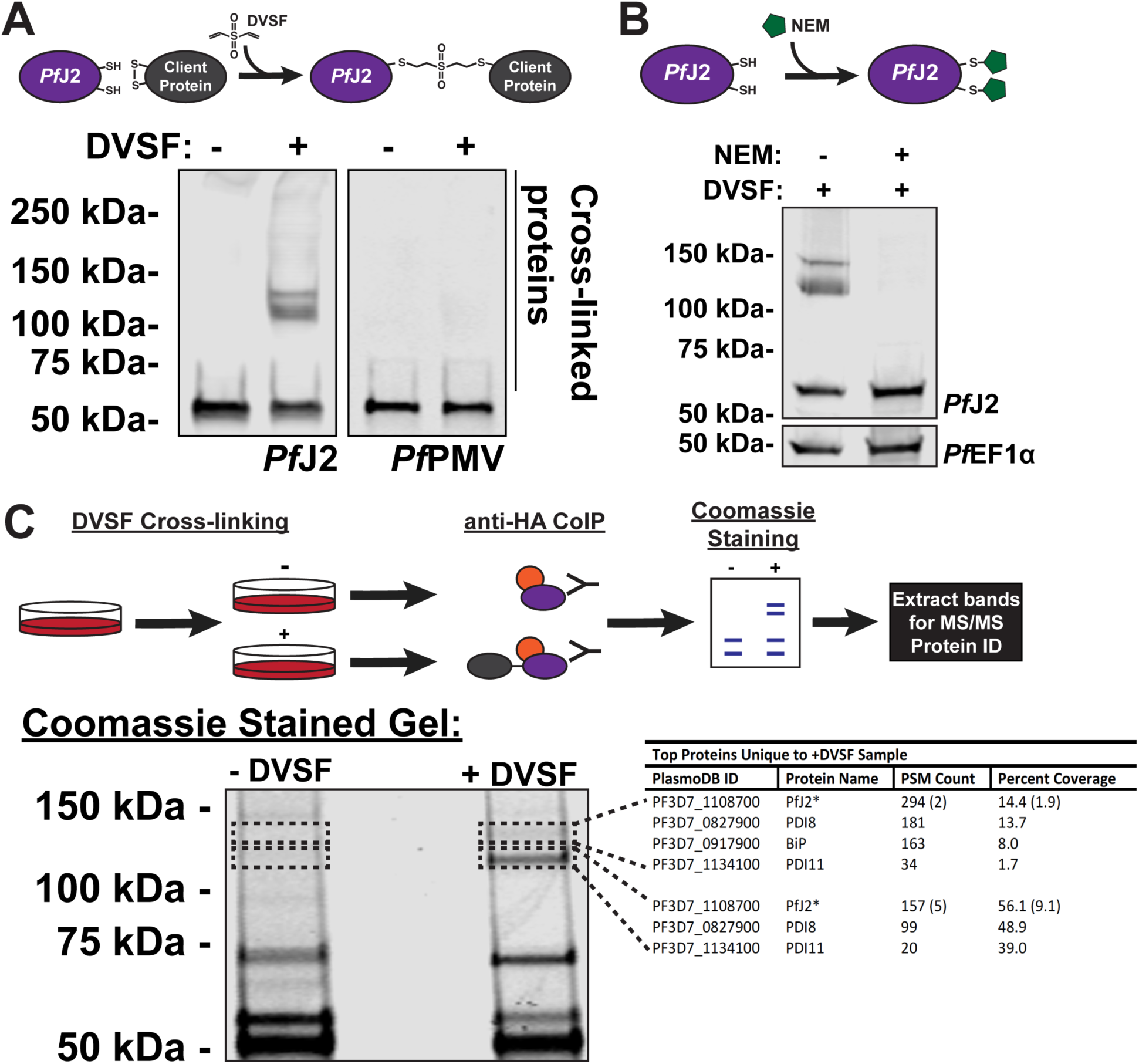
*Pf*J2 redox partners identified as *Pf*PDI8 and *Pf*PDI11. **A)** *Pf*J2^apt^ parasites were incubated with 3 mM divinyl sulfone (DVSF) in 1x PBS for 30 minutes at 37°C, then samples were taken for western blot analysis. Membranes were incubated with antibodies against HA and *Pf*PMV. **B)** *Pf*J2^apt^ parasite cultures were incubated with 1 mM N-ethylmaleimide (NEM) for 3 hours prior to removal of NEM and addition of 3 mM DVSF as described above. Samples were taken for western blot analysis. Membranes were incubated with antibodies against HA and *Pf*EF1α. **C)** *Pf*J2^apt^ parasite cultures were evenly split into two conditions: 3 mM DVSF or PBS only for 30 minutes at 37°C, after which parasite lysates were used for anti-HA immunoprecipitation. Immunoprecipitated proteins were separated via SDS-PAGE and visualized using Coomassie. Bands unique to the DVSF-treated sample were extracted, along with the corresponding section of gel in the untreated sample. Proteins were identified by tandem mass spectrometry, and proteins identified in both plus and minus DVSF samples eliminated for further study. The GeneID, protein name, PSM count, and percent coverage for all proteins with more than 1% coverage are shown in the table. A small amount of PfJ2 was identified in the -DVSF samples, and the PSM count and percent coverage is shown in parentheses.

### *Pf*PDI8 and *Pf*PDI11 are *Pf*J2 redox partners

Having shown the redox activity of *Pf*J2, we next sought to specifically identify those redox partnerships. To identify the proteins trapped to *Pf*J2 by DVSF, cultures were treated with the compound and immunoprecipitation of *Pf*J2 was performed (Figure 3C). As a control, the immunoprecipitation was also performed in parallel using cultures that had not received DVSF treatment. The immunoprecipitated proteins were subjected to separation by SDS-PAGE and visualized using Coomassie (Figure 3C). Two bands between 100-150 kDa, corresponding to those previously detected by western blot, were extracted from the +DVSF sample, along with the corresponding areas of the gel in the untreated samples (Figure 3C, perforated boxes). Proteins present in these gel slices were identified via MS/MS analysis. By analyzing the untreated control samples, we were able to remove background proteins and found that the top gel slice primarily contained *Pf*J2, *Pf*PDI8, *Pf*PDI11 and *Pf*BiP, and the bottom slice contained *Pf*J2, *Pf*PDI8, and *Pf*PDI11 (Figure 3C, table). The complete list of proteins identified in all samples can be found in Supplementary Table 2. Because alterations to *Pf*BiP migration during SDS-PAGE after DVSF treatment were not detected (Supplementary Figure 3), we chose to focus our attentions on *Pf*PDI8 and −11.

### *Pf*PDI8 and *Pf*PDI11 are redox-active ER proteins

Like *Pf*J2, *Pf*PDI8 and −11 are both predicted members of the Trx superfamily (Figure 4A). PfPDI8 appears to be a canonical PDI, with two Trx domains containing CXXC active sites that likely allow it to carry out disulfide oxidoreductase/isomerase activity. PfPDI11 also has two Trx domains, but each contains an unusual CXXS active site. *Pf*PDI8 has been characterized recombinantly *in vitro*, but both *Pf*PDI8 and −11 remain unstudied *in vivo* (Mahajan et al., 2006; Mouray et al., 2007). In order to validate interaction between these PDIs and *Pf*J2, and to understand their roles in the *P. falciparum* ER, we used the *glmS* ribozyme to create conditional knockdown parasite lines for each protein in the background of *Pf*J2^apt^ parasites (Prommana et al., 2013) (Figure 4B). Using CRISPR/Cas9 genome editing, we introduced sequences for a 3xV5 tag and the *glmS* ribozyme into the *pfpdi8* or the *pfpdi11* locus (*Pf*J2^apt^-PDI8^glmS^ and *Pf*J2^apt^-PDI11^glmS^, respectively) (Figure 4C). Correct modifications of the loci were validated by PCR integration test, and V5-tagged proteins were visualized by western blot (Figure 4D, E).

**Figure 4.**
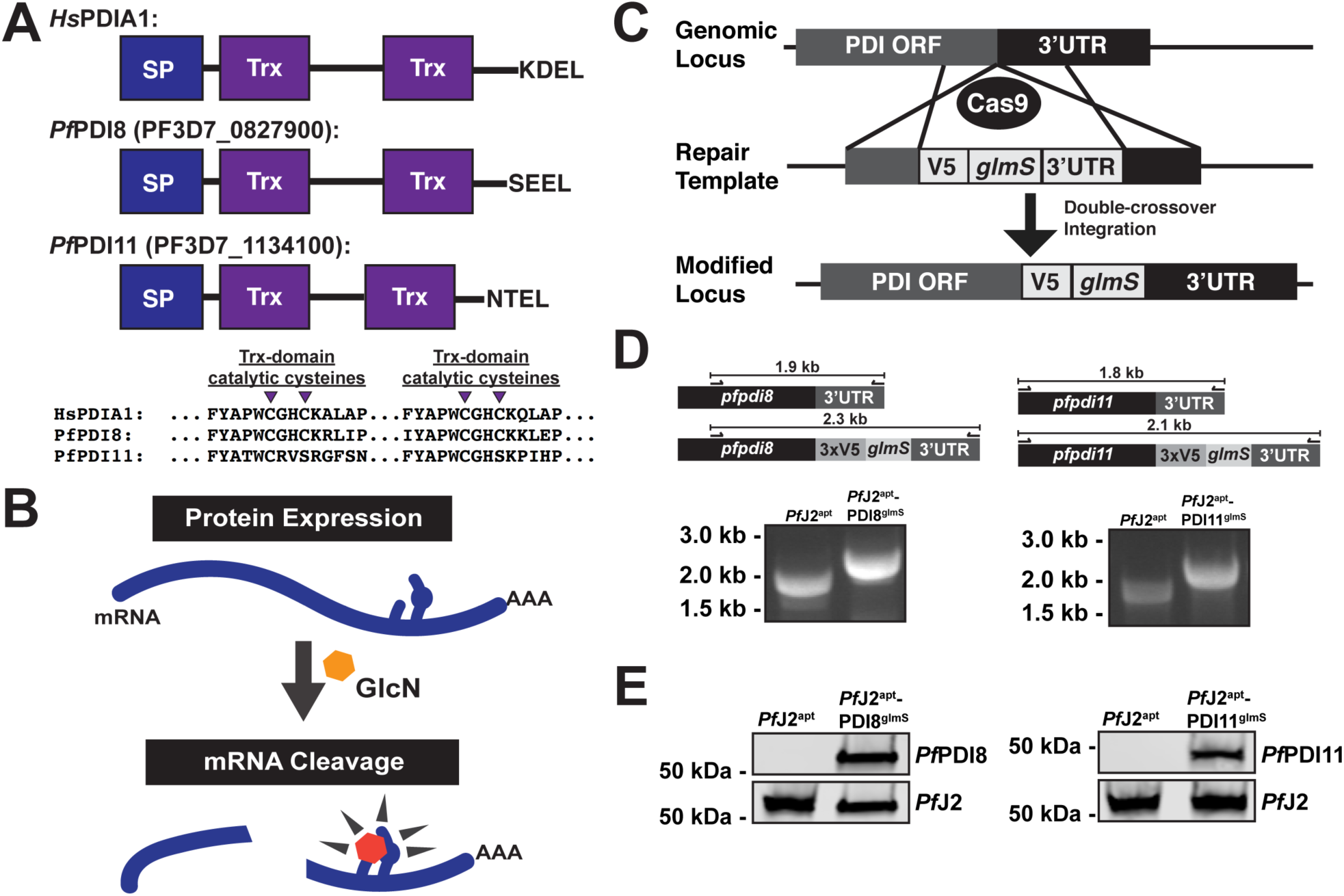
Generation of *Pf*PDI8 (PF3D7_0827900) and *Pf*PDI11 (PF3D7_1134100) conditional knockdown mutants using CRISPR/Cas9. **A)** Predicted domain structure of *Pf*PDI8, *Pf*PDI11, and human PDIA1, showing signal peptide (SP), thioredoxin domains (Trx), and C-terminal ER retention signals. Essential, conserved cysteine residues are shown for each of the proteins’ Trx domains. **B)** Regulation of protein expression using the *glmS* ribozyme system. The mRNA of interest encodes the ribozyme in the 3’UTR. Upon addition of glucosamine (GlcN, orange hexagon), which is converted to glucosamine-6-phosphate (pink hexagon) by the parasite, the ribozyme is activated to cleave the mRNA, leading to transcript instability and degradation (Prommana et al., 2013) **C)** Schematic of CRISPR/Cas9 mediated introduction of the *glmS* knockdown system into the genome. A repair template was transfected, along with a plasmid to express Cas9 and a gRNA, to introduce sequences for a 3xV5 tag, ER retention signals, stop codon, and *glmS* ribozyme. **D)** PCR integration test confirming correct modification of *pfpdi8* and *pfpdi11*. Correct integration results in increased amplicon size due to the V5 and *glmS* sequences. **E)** Western blots showing V5-tagged proteins in the *Pf*J2^apt^-PDI8^glmS^ and *Pf*J2^apt^-PDI11^glmS^ parasite lines at the predicted sizes for *Pf*PDI8 and −11.

The subcellular localizations of *Pf*PDI8 and −11 were determined by IFA, and both proteins were found to co-localize with *Pf*J2 in the ER (Figure 5A). To test the functionality of the *glmS* ribozyme knockdown system, each parasite line was treated with glucosamine (GlcN), and samples were taken for western blot analysis over the course of the parasite lifecycle. Compared to -GlcN control samples, protein levels were found to be reduced during GlcN treatment (Supplementary Figure 4). In order to determine the effect that PDI knockdown had on parasite growth, each cell line was treated with GlcN and parasite growth was measured over the course of two life cycles. We observed dramatic inhibition of parasite growth during *Pf*PDI8 knockdown, but no growth defects were observed when *Pf*PDI11 was knocked down (Figure 5B). These results demonstrate that *Pf*PDI8 is essential for the asexual lifecycle and suggest that *Pf*PDI11 may be dispensable.

**Figure 5.**
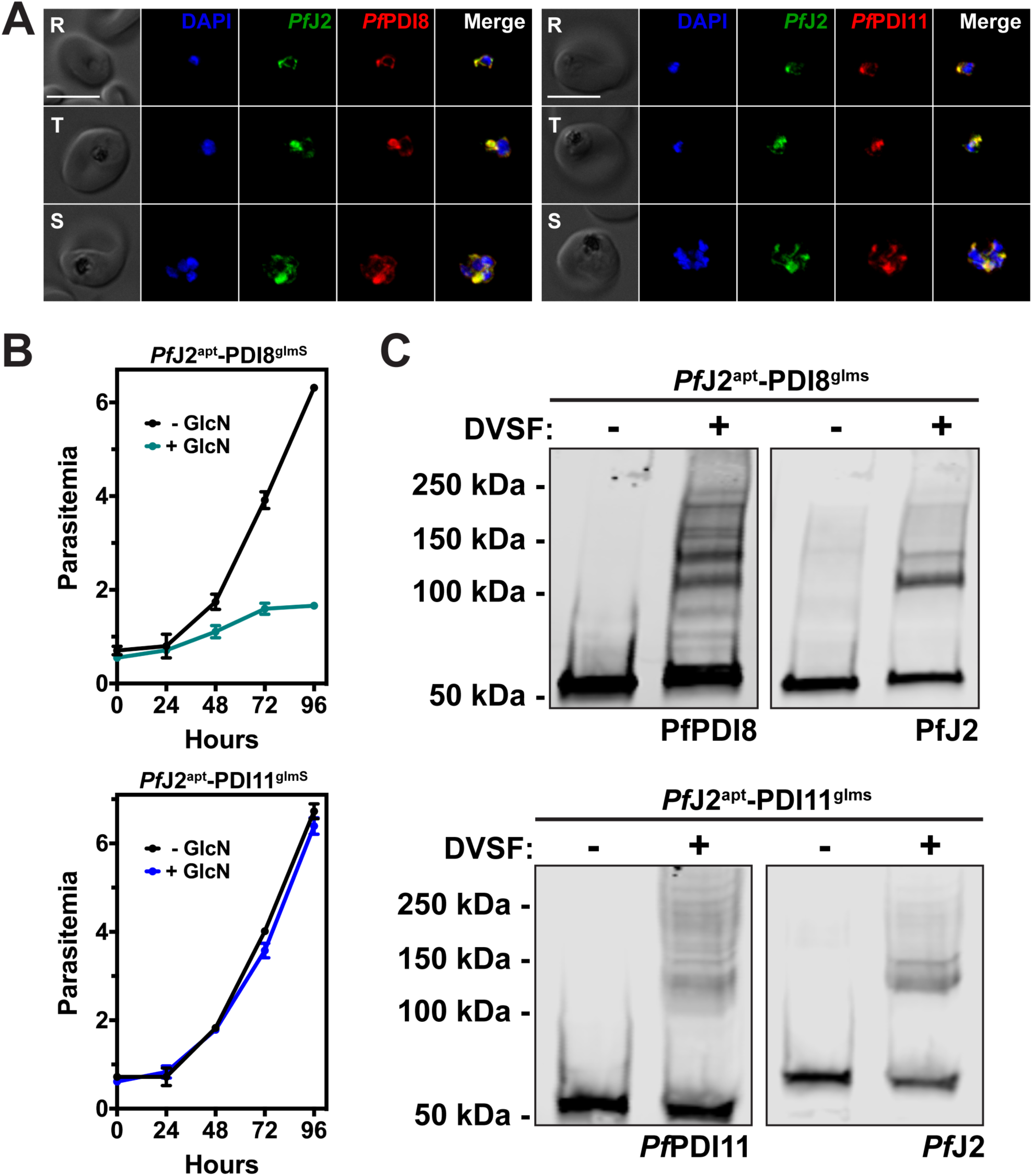
*Pf*PDI8 and *Pf*PDI11 are redox-active ER proteins. **A)** *Pf*J2^apt^-PDI8^glmS^ and *Pf*J2^apt^-PDI11^glmS^ parasites were fixed with paraformaldehyde and glutaraldehyde and stained with DAPI (blue) and with antibodies against HA (green) and V5 (red). Ring (R), Trophozoite (T), and Schizont (S) stage parasites are shown. Z-stack Images were deconvoluted and shown as a single, maximum intensity projection. Scale bar represents 5 μm. **B)** Asynchronous *Pf*J2^apt^-PDI8^glmS^ (top) and *Pf*J2^apt^-PDI11^glmS^ (bottom) parasites were grown in normal (-GlcN) or knockdown (+ 5mM GlcN) conditions, and parasite growth was monitored daily for 96 hours via flow cytometry. **C)** *Pf*J2^apt^-PDI8^glmS^ and *Pf*J2^apt^-PDI11^glmS^ parasites were incubated with 3 mM DVSF in 1x PBS for 30 minutes at 37°C, then samples were taken for western blot analysis. Membranes were incubated with antibodies against HA (*Pf*J2) and V5 (*Pf*PDI8 or −11).

*Pf*PDI8 contains two thioredoxin domains with classical CXXC active site cysteines, allowing the protein to function in oxidative folding as an oxidoreductase/isomerase (Mahajan et al., 2006; Mouray et al., 2007). *Pf*PDI11 has two thioredoxin domains containing noncanonical CXXS active sites, but likely maintains the ability to form mixed disulfide bonds with client proteins through the conserved cysteine residues (Anelli, 2003; Fomenko & Gladyshev, 2009; Park et al., 2009). To determine whether we could trap redox interactions between these PDIs and their substrates, *Pf*J2^apt^-PDI8^glmS^ and *Pf*J2^apt^-PDI11^glmS^ cultures were treated with DVSF, and protein lysates were collected for western blot analysis. Several high molecular weight bands containing *Pf*PDI8 appear following DVSF treatment, indicating that multiple substrates rely on the redox activity of *Pf*PDI8, in contrast to *Pf*J2, whose western blot shows a narrower set of redox substrates (Figure 5C). Similar results were observed for PfPDI11 (Figure 5C).

We next sought to determine if DVSF crosslinking occurs through Trx-domain cysteines. To do this, we attempted to generate parasites overexpressing either wild-type copies of *Pf*J2, *Pf*PDI8, *Pf*PDI11, or overexpressing these proteins with cysteine-to-alanine mutations in the Trx domain active sites. These types of mutations abolish DVSF-crosslinking in Trx proteins of model organisms (Araki et al., 2017; Naticchia et al., 2013). We were unable to generate parasites overexpressing wild-type or mutant copies of *Pf*J2. We were also unable to generate parasites overexpressing a mutant version of *Pf*PDI8, but were successful in creating parasites overexpressing the wild-type protein; characterization of that parasite line revealed mislocalization of the overexpressed *Pf*PDI8 (Supplementary Figure 5). Given the essential nature of *Pf*J2 and *Pf*PDI8, we concluded that the parasites may be sensitive to their overexpression and to mutations in their Trx domains.

In contrast, we were successfully able to generate both wild-type and cysteine-to-alanine *Pf*PDI11 overexpression mutants (*Pf*PDI11^*wt*^ and *Pf*PDI11^*mut*^, respectively) (Supplementary Figure 6). Both parasite lines displayed the expected ER co-localization with PfJ2 (Supplementary Figure 6). Importantly, treatment of these parasites with DVSF revealed extensive crosslinking between *Pf*PDI11 and substrates in the wild-type parasites, but crosslinking is abolished in parasites with cysteine-to-alanine mutations in the Trx domain (Supplementary Figure 6). These data demonstrated the specificity of DVSF for trapping redox partnerships in *P. falciparum*.

Given the unusual nature of the *Pf*PDI11 CXXS Trx-domain active site, we took advantage of these overexpression parasites to further investigate *Pf*PDI11 function. Trx-domain-proteins with CXXS active sites are largely under-studied in all organisms (Anelli, 2003; Fomenko & Gladyshev, 2009; Park et al., 2009). It is possible that the remaining active-site cysteines in *Pf*PDI11 forms mixed disulfides that may serve to retain proteins the in ER, prevent their aggregation, and/or block cysteines from non-productive bond formation as they fold (Anelli, 2003; Park et al., 2009). Therefore, we hypothesized that we would be able to detect the mixed *Pf*PDI11-substrate disulfide bonds by non-reducing SDS-PAGE and western blotting. Indeed, we were able to detect high-molecular-weight species of *Pf*PDI11 when *Pf*PDI11^wt^ parasite lysates were used for western blotting under non-reducing conditions (Supplementary Figure 7). In contrast, these species were missing when *Pf*PDI11^mut^ parasite lysates were used (Supplementary Figure 7).

### The *Pf*BiP-*Pf*J2-*Pf*PDI8 oxidative folding complex

Having shown that *Pf*J2 and *Pf*PDI8 are redox partners and that both proteins are essential for the *P. falciparum* asexual lifecycle, we decided to focus on their interaction and what roles they may play together in the ER. To confirm the interaction between *Pf*J2 and *Pf*PDI8, *Pf*J2 was immunoprecipitated from *Pf*J2^apt^-PDI8^glmS^ parasites, and *Pf*PDI8 was found to co-immunoprecipitate (Figure 6A). When performing the reciprocal co-IP, we were unable to detect *Pf*J2 pulling down with *Pf*PDI8, perhaps due to inefficiency of the anti-V5 IP (Supplemental Figure 8). However, we were able to detect a band of overlapping *Pf*J2/*Pf*PDI8 signal when the *Pf*PDI8 IP was performed on cultures treated with DVSF, showing that the two proteins are interacting, redox partners (Figure 6B).

**Figure 6.**
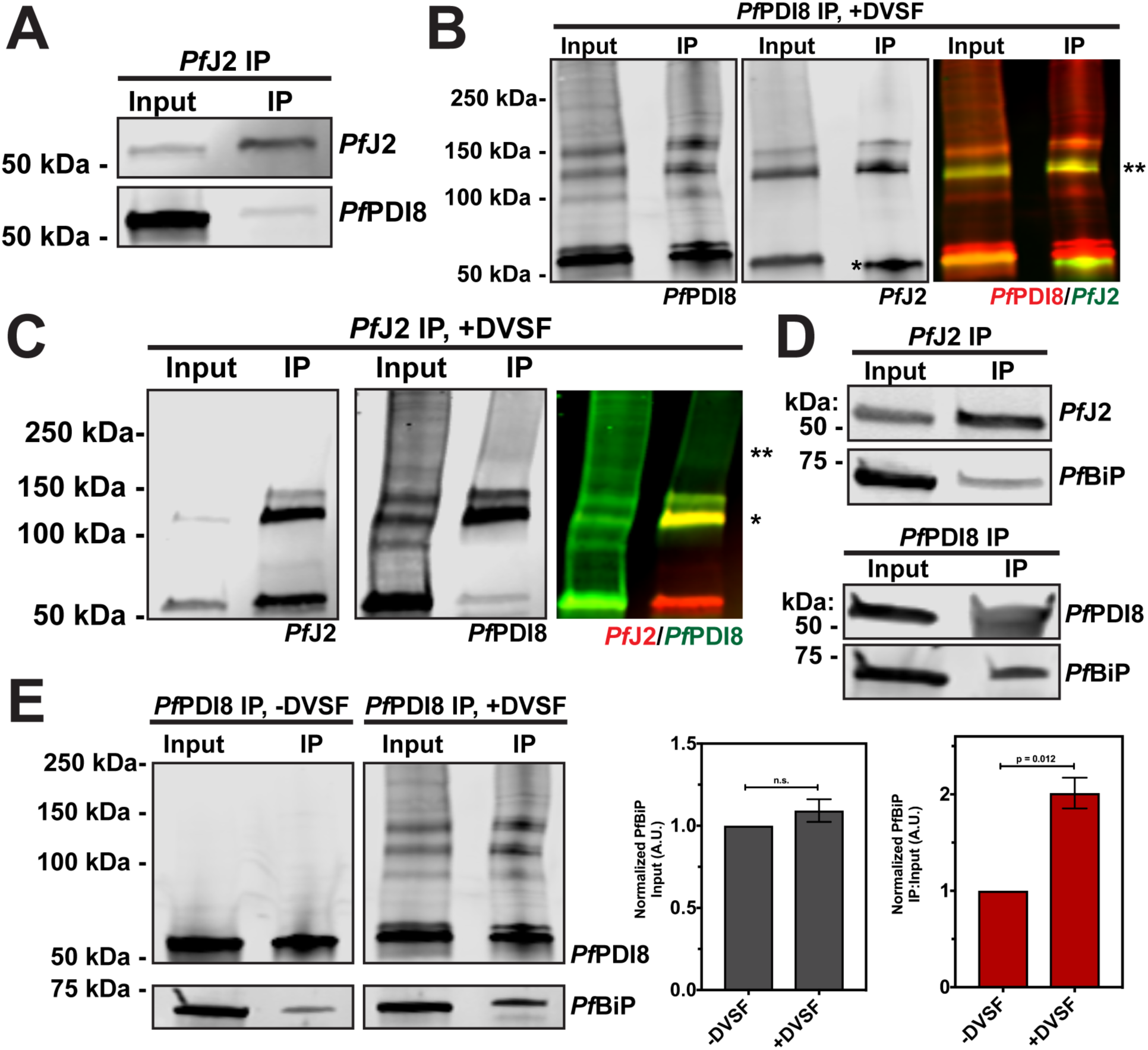
The *Pf*BiP-*Pf*J2-*Pf*PDI8 oxidative folding complex. **A)** *Pf*J2 and interacting proteins were immunoprecipitated from *Pf*J2^apt^-PDI8^glmS^ parasite lysate using anti-HA antibodies. Input and eluted IP samples were used for western blot analysis. Membrane was probed with HA and V5 antibodies to detect *Pf*J2 and *Pf*PDI8, respectively. **B)** *Pf*J2^apt^-PDI8^glmS^ parasites were incubated with 3 mM DVSF as described above, then V5 antibodies were used to immunoprecipitate *Pf*PDI8 and interacting proteins. Input and eluted IP samples were used for western blot analysis. Membrane was probed with V5 and HA antibodies to detect *Pf*PDI8 and *Pf*J2, respectively. Antibody heavy chain is indicated by the single asterisk (*) in the *Pf*J2 panel. A merged image of the *Pf*PDI8 (red) and *Pf*J2 (green) signal is shown, with the yellow overlap in signal indicated by a double asterisk (**). **C)** *Pf*J2^apt^-PDI8^glmS^ parasites were incubated with 3 mM DVSF as described above, then HA antibodies were used to immunoprecipitate *Pf*J2 and interacting proteins. Input and eluted IP samples were used for western blot analysis. Membrane was probed with HA and V5 antibodies to detect *Pf*J2 and *Pf*PDI8, respectively. A merged image of the *Pf*J2 (red) and *Pf*PDI8 (green) signal is shown, with a single asterisk (*) indicating the yellow overlap in signal and a double asterisk (**) indicating *Pf*PDI8+subtrates that co-immunoprecipitated with *Pf*J2. **D)** *Pf*J2 and *Pf*PDI8 were immunoprecipitated from *Pf*J2^apt^ and *Pf*J2^apt^-PDI8^glmS^ parasite lysates, respectively. Input samples and eluted IP proteins were used for western blot analysis. Membrane was probed with HA and *Pf*BiP antibodies (top) or V5 and *Pf*BiP antibodies (bottom). **E)** *Pf*J2^apt-^PDI8^glmS^ parasite cultures were evenly split into two conditions: 3 mM DVSF or PBS only for 30 minutes at 37°C, after which parasite lysates were used for anti-V5 immunoprecipitation. Input and eluted IP proteins were analyzed by western blot using V5 and *Pf*BiP antibodies. The PfBiP signal was measured for each lane and the ratio of IP-to-Input signal was determined. N = 3 biological replicates.

Our observations using the redox crosslinker DVSF showed that *Pf*PDI8 is a major redox partner for *Pf*J2, whereas *Pf*PDI8 has multiple other redox partnerships (Figures 3C, 5D). One explanation for this observation is that *Pf*J2 may work upstream to prime *Pf*PDI8 for interaction with its substrates. Therefore, we asked whether we could detect *Pf*PDI8+substrates co-immunoprecipitating with *Pf*J2. When *Pf*J2 was immunoprecipitated from *Pf*J2^apt^-PDI8^glmS^ parasites treated with DVSF, we were able to detect *Pf*PDI8 trapped to other substrates, indicated by smearing of the *Pf*PDI8 signal above 150 kDa (Figure 6C). Together, these results confirm the interaction between *Pf*J2 and *Pf*PDI8 and suggest that *Pf*J2 may be part of a complex including *Pf*PDI8 and its substrates.

Oxidative folding in the ER, mediated by proteins such as *Pf*J2 and *Pf*PDI8, is only one aspect of protein folding in the ER, and likely works in conjunction with other folding determinants, such as the Hsp70 BiP. *Pf*J2 is an ER Hsp40—a class of co-chaperones that interact with the Hsp70 BiP—and BiP is likely involved in the folding of the same substrates that interact with *Pf*PDI8. Such interactions between oxidative folding chaperones and BiP likely exist in the ER of other organisms, but it is not known how these types of proteins might work together to ensure substrates reach their native states. Given the smaller repertoire of ER Trx-superfamily proteins present in *P. falciparum*, the parasite is an ideal choice for investigating the relationship between these proteins and BiP. Therefore, we next asked whether *Pf*BiP interacts with *Pf*J2 and/or *Pf*PDI8.

Western blot analysis of proteins co-immunoprecipitating with *Pf*J2 revealed that *Pf*BiP does interact with *Pf*J2, consistent with our *Pf*J2 co-IP experiments and the proteins’ predicted chaperone/co-chaperone roles (Table 1, Figure 6D, top). Lack of a suitable antibody precluded reciprocal *Pf*BiP immunoprecipitation to probe for *Pf*J2. However, as a control we showed that *Pf*BiP is not detected when wild-type parasites (lacking HA-tagged *Pf*J2) are subjected to anti-HA immunoprecipitation, ruling out nonspecific binding during the co-IP experiment (Supplementary Fig 9).

Next, we found that when *Pf*PDI8 was immunoprecipitated, *Pf*BiP was detected (Figure 6D, bottom). We further reasoned that because the same substrates may rely on both *Pf*PDI8 and *Pf*BiP to achieve their native state, trapping the *Pf*PDI8-substrate interaction with DVSF may increase the amount of *Pf*BiP co-immunoprecipitating with *Pf*PDI8. To test this hypothesis, parasite cultures were equally split into +/- DVSF treated aliquots, *Pf*PDI8 was immunoprecipitated, and lysates probed for *Pf*PDI8 and *Pf*BiP. Consistent with our hypothesis, we detected a two-fold increase in the amount of *Pf*BiP that pulled down with *Pf*PDI8 crosslinked to its substrates, with no significant difference in the starting amount of *Pf*BiP detected in the sample prior to immunoprecipitation (Figure 6E). These results suggest that *Pf*J2 and *Pf*PDI8 work together with the major ER folding chaperone *Pf*BiP to help substrates reach their native states.

### ER redox interactions are druggable

Our data have shown that *Pf*J2 and *Pf*PDI8, which participate in ER redox partnerships with each other as well as other substrates, are essential proteins in the *P. falciparum* asexual lifecycle. These proteins, through inhibition of their redox interactions, may represent unexploited targets for antimalarials. Further, our data show that DVSF can target these redox partners, suggesting that these proteins may be druggable. Indeed, consistent with the idea of targeting ER redox proteins in disease, high-throughput drug screens have identified potent inhibitors of human PDI in an effort to combat upregulation that is associated with some cancers and neurodegenerative diseases (Hoffstrom et al., 2010; Kaplan et al., 2015; Vatolin et al., 2016; Xu et al., 2012).

We tested four of these commercially available PDI inhibitors—16F16, LOC14, CCF642, and PACMA31—for activity against cultured asexual *P. falciparum* parasites. In contrast to their reported, highly potent activity against human cells, we observed a wide range of IC_50_ values for *P. falciparum* (Supplementary Fig 10). The compound with the best anti-*Plasmodium* activity was 16F16, with an IC_50_ value of approximately 4 μM (Figure 7A) (Harbut et al., 2012).

**Figure 7.**
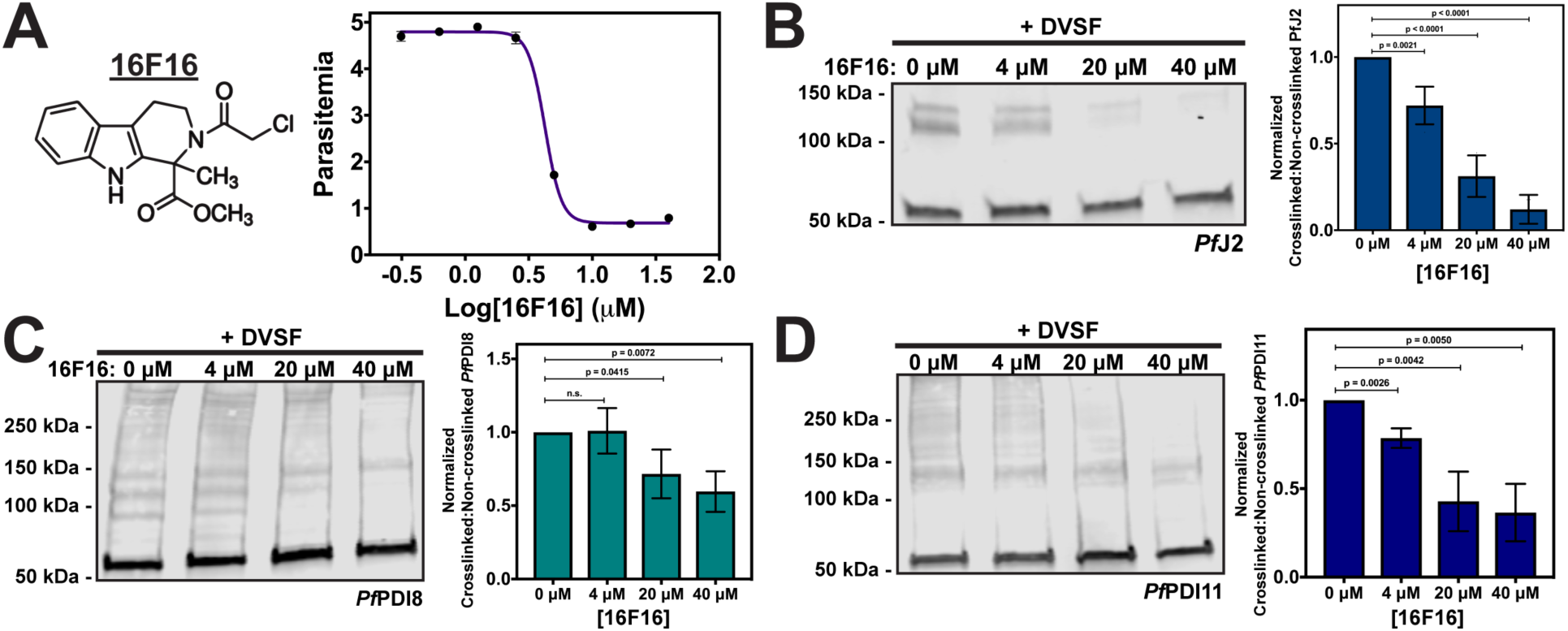
ER redox interactions are sensitive to interruption by a small molecule. Asynchronous *Pf*J2^apt^ parasites were incubated in various concentrations of the human PDI inhibitor 16F16. Parasite growth was determined via flow cytometry at 72 hours and the 16F16 IC_50_ was determined to be approximately 4 μM. Each data point in the curve represents the mean parasitemia at a given concentration, in technical triplicate. Error bars, which are not shown for data points in which they are smaller than the circle symbol, represent standard deviation from the mean. **B)** *Pf*J2^apt^, **C)** *Pf*J2^apt^-PDI8^glmS^, and **D)** *Pf*J2^apt^-PDI11^glmS^ parasites cultures were equally split and incubated with three concentrations of 16F16 for 3 hours prior to removal of 16F16, then incubation with 3 mM DVSF as described above. Samples were taken for western blot analysis, loading equal parasite equivalents into each gel. Membranes were incubated with antibodies against HA or V5. Signal for non-crosslinked (the band at approximately 50 kDa) and crosslinked proteins (>50 kDa) was measured. Inhibition was measured by determining the ratio of crosslinked to non-crosslinked signal. N = 3 biological replicates for each parasite line.

16F16 inhibits human PDI function by covalently binding the cysteines of the Trx domain active sites, thereby blocking their ability to catalyze oxidative folding (Hoffstrom et al., 2010; Kaplan et al., 2015). If 16F16 behaves similarly in *P. falciparum*, we reasoned that treatment of cultures with 16F16 prior to performing redox crosslinking with DVSF would prevent crosslinking from occurring, assuming that both compounds rely on the same redox-active cysteine residues for their activity. Indeed, pre-treatment with increasing amounts of 16F16 significantly and reproducibly decreased the amount of crosslinked *Pf*J2 detected by western blot (Figure 7B). Our data indicate that after DVSF treatment, the high molecular weight bands detected consist of *Pf*J2, *Pf*PDI8, and *Pf*PDI11 (Figure 3). Therefore, the observed reduction in *Pf*J2 crosslinking likely occurs due to direct reaction of 16F16 with the *Pf*J2 Trx-domain active site and/or the *Pf*PDI8 and −11 active sites. Similar experiments showed that pre-treatment of cultures with 16F16 also blocked crosslinking of *Pf*PDI8 and −11 with their substrates, though to a lesser extent than what was observed for PfJ2 (Figure 7C, D).

These data demonstrate that redox interactions within the *P. falciparum* ER, occurring between essential proteins like *Pf*J2 and *Pf*PDI8 and their substrates, are sensitive to small molecule inhibition. Additionally, the disparity in activity observed for the PDI inhibitors against human and *P. falciparum* cell lines suggest that development of *Plasmodium*-specific inhibitors is likely possible (Supplementary Figure 10).

## Discussion

The ability to conduct oxidative folding likely underlies the diverse functions of the *P. falciparum* ER. The oxidizing environment of the ER encourages disulfide bond formation, but only the correct bonds allow proteins to reach their native states. Therefore, organisms must maintain a way to reduce/isomerize nonproductive disulfides. We have used CRISPR/Cas9 genome editing and conditional knockdown to show here that a putative disulfide reductase in the *P. falciparum* ER—*Pf*J2—Is essential for the parasite asexual lifecycle (Figures 1, 2).

A co-IP/mass spectroscopy approach with stringent parameters for identifying interacting partners places *Pf*J2 in the broader context of ER biology, revealing that *Pf*J2 interacts with other folding determinants, such as BiP and Endoplasmin, as well as other members of the Thioredoxin superfamily, such as PDIs. The remaining proteins that were identified, most with unknown localization throughout the secretory pathway and many with no known function, may represent substrates that rely on *Pf*J2 and these other chaperones for their folding and/or trafficking. Among the proteins identified were large, complex proteins such as *Pf*MSP1 and *Pf*RhopH3 (Table 1). Both proteins have numerous cysteine residues that must navigate oxidative folding when they are synthesized into the ER, likely relying on *Pf*J2 and *Pf*PDIs to do so correctly. Consistent with this hypothesis, a recent study identified *Pf*J2, *Pf*PDI11, and *Pf*Endoplasmin as potential contributors to folding and trafficking of *Pf*EMP1, a cysteine-rich transmembrane protein that serves as the major *P. falciparum* virulence factor (Batinovic et al., 2017).

One major *Pf*J2 redox substrate—*Pf*PDI8—was identified using a chemical biology approach. DVSF is a redox-specific crosslinker that has been used to identify redox partnerships between Thioredoxin proteins in the cytoplasm of model organisms (Allan et al., 2016; Araki et al., 2017; Naticchia et al., 2013). To our knowledge, this compound had not yet been used to trap and define redox partnerships in *Plasmodium*, nor in the ER of any organism. We demonstrate its utility in the *Plasmodium* ER, using it to identify the redox partnership between *Pf*J2, *Pf*PDI8 and *Pf*PDI11(Figure 3). Double conditional *P. falciparum* mutants showed that *Pf*PDI8 is also an essential, ER-resident protein and allowed us to probe more deeply the relationship between *Pf*J2 and *Pf*PDI8 (Figures 4-6). *Pf*J2 likely acts as a reductase, and previous *in vitro* characterization of *Pf*PDI8 revealed that it behaves like a classical PDI, capable of both forming and reducing disulfide bonds (Cunnea et al., 2003; Mahajan et al., 2006; Mouray et al., 2007; Oka et al., 2013; Ushioda et al., 2008).The propensity for *Pf*PDI8 to use its Trx domains either for oxidation of cysteines or reduction of disulfides likely depends on the oxidation state of its own active site (i.e. reduced *Pf*PDI8 can act as a reductase). One explanation for the redox partnership between *Pf*J2 and *Pf*PDI8 is that *Pf*J2 primes *Pf*PDI8 to act as a reductase for some or all of the numerous substrates we visualized using DVSF (Figure 5D). Consistent with this hypothesis, immunoprecipitation experiments showed that *Pf*PDI8+substrates pull down with *Pf*J2 (Figure 6C). We also found that *Pf*J2 and *Pf*PDI8 both interact with the Hsp70 *Pf*BiP, and detection of that interaction increases for *Pf*PDI8 when it is trapped to its substrates (Figure 6D, E). These data suggest a model in which *Pf*J2, *Pf*PDI8, and *Pf*BiP cooperate to ensure substrates in the ER correctly navigate the oxidative folding process to achieve their native states (Figure 8).

**Figure 8.**
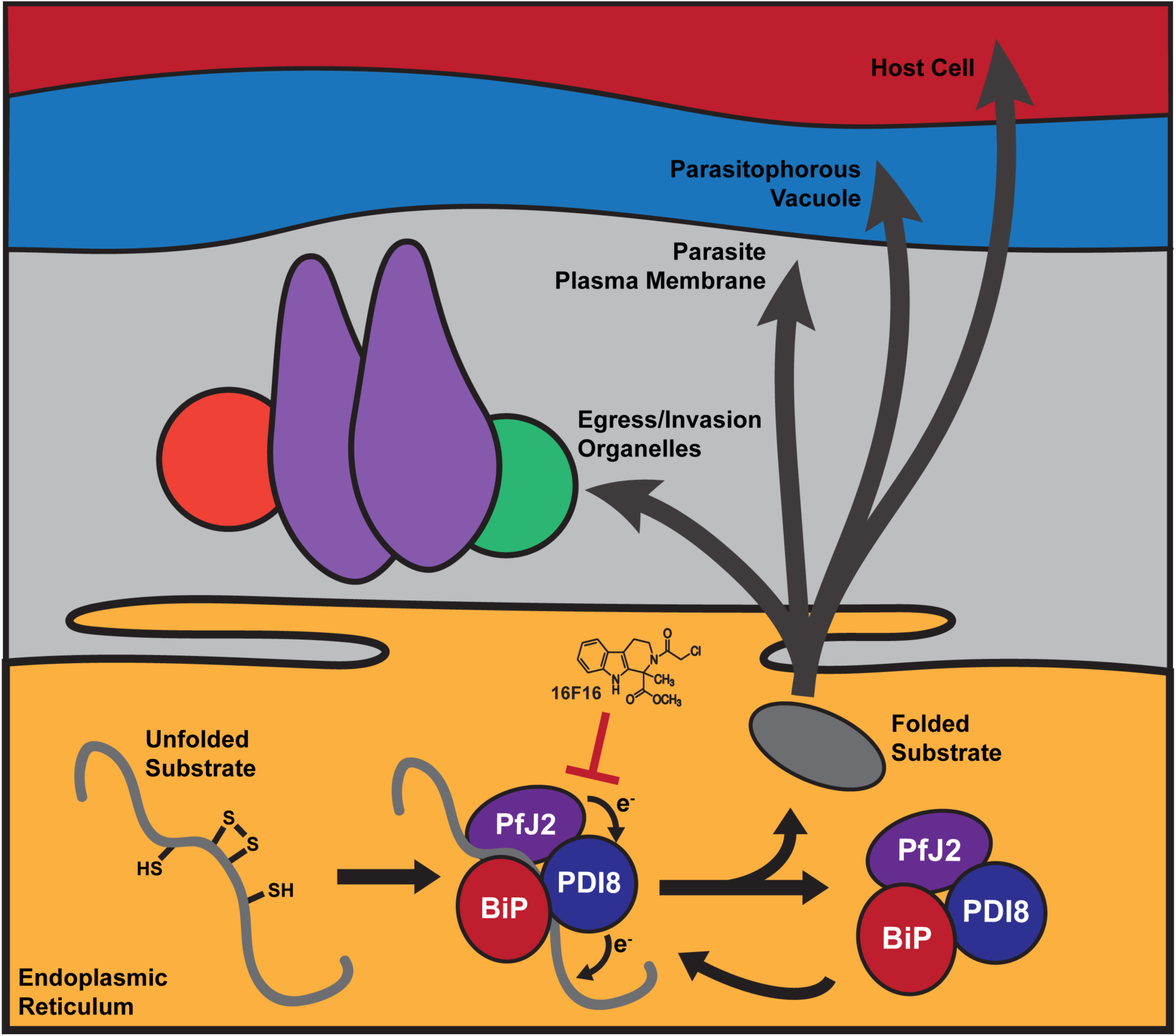
Oxidative folding in the *P. falciparum* ER. We propose that Trx-domain proteins like *Pf*J2 and *Pf*PDI8 work with *Pf*BiP to help nascent proteins, which perform essential functions within the ER and throughout the parasite secretory pathway, achieve their native states. The redox interactions between *Pf*J2, *Pf*PDI8, and their substrates are sensitive to inhibition by small molecules like 16F16, which could be expected to disrupt oxidative folding and impair the parasite’s ability to perform functions essential for survival and replication.

We also identified *Pf*PDI11 as a redox substrate of *Pf*J2 (Figure 3). Our data demonstrate that *Pf*PDI11 retains the ability to form mixed disulfides with client proteins despite the unusual CXXS Trx-domain active site (Figure 5, Supplementary Figures 6, 7). Typically, the second cysteine of the Trx-domain active site is used to resolve enzyme-substrate mixed disulfides (Hatahet & Ruddock, 2009). Therefore, the mechanisms used to resolve mixed disulfides between CXXS active sites and their substrates remains unclear, both in *P. falciparum* and other organisms. We propose that an ER-resident reductase such as *Pf*J2 helps resolve mixed disulfides, which would explain why *Pf*J2 and *Pf*PDI11 were found to be redox partners.

Importantly, given the recent stagnation observed in malaria elimination efforts, which is coincident with increasing cases of antimalarial resistance, we not only identified two proteins with essential functions; we further demonstrated that the redox partnerships of these proteins are sensitive to disruption by small molecule inhibition (Figure 7). 16F16 is a covalent inhibitor that blocks Trx-domain cysteines (Hoffstrom et al., 2010; Kaplan et al., 2015). Such a compound, if specific for *P. falciparum*, could be expected to cripple oxidative folding in the ER and kill the parasite. Recently, interest in covalent inhibitors for treatment of human disease has renewed, with several covalent inhibitors approved for use by the United States Food and Drug Administration (Ghosh et al., 2019). One particular concern with covalent inhibitors is the fact that mutagenesis of the target residue would result in resistance, but mutagenesis of Trx-domain cysteines would lead to loss of function in and of itself, presumably making this type of resistance harder to evolve. Finally, given the disparity in activity observed for the PDI inhibitors against human cell lines and *P. falciparum*, enough diversity likely exists between these conserved proteins that *Plasmodium*-specific inhibitors could be developed (Supplementary Figure 10). Therefore, essential Trx-domain proteins in the parasite ER— like *Pf*J2 and *Pf*PDI8—represent a class of proteins and a pathway in the ER that is apt for antimalarial drug development.

## Materials and Methods

### Construction of Plasmids

Parasite genomic DNA was isolate from 3D7 parasites using QIAamp DNA blood kit (QIAGEN). All constructs utilized in this study were confirmed by sequencing. Plasmids were constructed using the Sequence and Ligation Independent Cloning (SLIC) method. Plasmids to express Cas9 and gRNAs were constructed using pUF1-Cas9 as previously described (Cobb et al., 2017; Ghorbal et al., 2014; Kudyba et al., 2018). All primers used in this study are listed in Supplemental Table 3. pfpdi8 cDNA was prepared using TRIzol-extracted mRNA and reverse transcription with primer P20 (SuperScript III, Invitrogen). All restriction enzymes used in plasmid construction were purchased from New England Biolabs.

To generate pMG74-PfJ2, approximately 500 bp of the sequence encoding the *Pf*J2 C-terminus was amplified using primers P1 and P2, and approximately 500 bp from the pfj2 3’UTR were amplified using P3 and P4. The two amplicons were joined together via PCR sewing using P1 and P4, then inserted into pMG74 (Ganesan et al., 2016) digested with AflII and AatII. For expression of a *Pf*J2 gRNA, oligos P31 and P32 were inserted into pUF1-Cas9.

To generate pV5-glmS-PDI8, approximately 500 bp of the sequence encoding the *Pf*PDI8 C-terminus was amplified using primers P5 and P6. The 3x V5 tag was added to this amplicon via PCR sewing using a linearized plasmid encoding the 3xV5 sequence and primers P5 and P7. The glmS ribozyme sequence was amplified from pHA-glmS (Prommana et al., 2013) using P8 and P9, then added to the *Pf*PDI8 C-terminus+V5 amplicon via PCR sewing using P5 and P9. The resulting amplicon was inserted into pHA-glmS that had been digested with AfeI and NheI, creating pPDI8-Cterm. Approximately 500 bp of the *pfpdi8* 3’UTR was amplified using P10 and P11, then inserted into pPDI8-Cterm that had been digested with HindIII and NotI, creating pV5-glmS-PDI8. For expression of a *Pf*PDI8 gRNA, oligos P33 and P34 were inserted into pUF1-Cas9.

To generate pV5-glmS-PDI11, approximately 500 bp of the sequence encoding the *Pf*PDI11 C-terminus was amplified using primers P12 and P13. The 3x V5 tag was added to this amplicon via PCR sewing using a linearized plasmid encoding the 3xV5 sequence and primers P12 and P14. The glmS ribozyme sequence was amplified from pHA-glmS (Prommana et al., 2013) using 15 and P16, then added to the *Pf*PDI8 C-terminus+V5 amplicon via PCR sewing using P12 and P16. The resulting amplicon was inserted into pHA-glmS that had been digested with AfeI and NheI, creating pPDI11-Cterm. Approximately 500 bp of the *pfpdi11* 3’UTR was amplified using P17 and P18, then inserted into pPDI11-Cterm that had been digested with HindIII and NotI, creating pV5-glmS-PDI11. For expression of a *Pf*PDI11 gRNA, oligos P35 and P36 were inserted into pUF1-Cas9.

*Pf*PDI8 and *Pf*PDI11 overexpression was carried out by using CRISPR/Cas9 to insert the open reading frame (ORF) of the tagged genes into the *pfhsp110c* locus. pUC57-Hsp110, the repair plasmid targeting *pfhsp110*, includes the last 429 bp encoding the PfHsp110c (PF3D7_0708800) C-terminus, a 2A skip peptide sequence, sequences for various peptide tags, then the first 400 bp from the *pfhsp110c* 3’UTR. This plasmid was synthesized by GeneScript. For expression of a *Pf*Hsp110c gRNA, oligos P37 and P38 were inserted into pUF1-Cas9.

To generate pUC57-Hsp110-PDI8^wt^, the *Pf*PDI8 ORF was amplified from cDNA using P19 and P20. A sequence encoding the 3xV5 was attached to this amplicon via PCR sewing using a linearized plasmid encoding the tag and primers P19 and P21. The resulting amplicon was inserted into pUC57-Hsp110 digested with MfeI and SpeI.

To generate pUC57-Hsp110-PDI11^wt^, the *Pf*PDI11 ORF was amplified using P22 and P23. A sequence encoding the 3xV5 was attached to this amplicon via PCR sewing using a linearized plasmid encoding the tag and primers P22 and P24. The resulting amplicon was inserted into pUC57-Hsp110 digested with MfeI and SpeI.

To generate pUC57-Hsp110-PDI11^mut^, which required mutagenesis of the 2 Trx-domain active site, the PfPDI11 ORF was amplified in 3 parts. Part 1 was amplified using P25 and P26. Part 2 was amplified using P27 and P28. Part 3 was amplified using P29 and P30. Parts 1+2 were joined together using PCR sewing and primers P25 and P28. The resulting amplicon was attached to Part 3 using PCR sewing and primers P25 and P30. A sequence encoding the 3xV5 was attached to this amplicon via PCR sewing using a linearized plasmid encoding the tag and primers P25 and P24. The resulting amplicon was inserted into pUC57-Hsp110 digested with MfeI and SpeI.

### Parasite Culture and Transfection

P. falciparum asexual parasites were cultured in RPMI 1640 medium supplemented with AlbuMAX I (Gibco) and transfected as described earlier (Drew et al., 2008; I. Russo et al., 2009).

To generate the *Pf*J2^apt^ parasite line, RBCs were transfected with 20 µg pMG74-PfJ2 (linearized prior to transfection using EcoRV) and 50 µg pUF1-Cas9-PfJ2, then fed to 3D7 parasites. Drug pressure was applied 48 hours after transfection, selecting for integration using 0.5 µM aTc and 2.5 µg/mL Blasticidin. After parasites grew back up from transfection and were cloned using limiting dilution, clones were maintained in medium containing 10 nM aTc and 2.5 µg/mL Blasticidin. Unless started otherwise, all +/- aTc growth experiments were conducted in medium containing 10 nM aTc and 2.5 µg/mL Blasticidin or medium containing only 2.5 µg/mL Blasticidin.

To generate the *Pf*J2^apt^-PDI8^glms^ parasite line, RBCs were transfected with 50 µg pV5-glmS-PDI8 and 50 µg pUF1-Cas9-PDI8, then fed to *Pf*J2^apt^ parasites. Drug pressure was applied 48 hours after transfection, selecting with 0.5 µM aTc, 2.5 µg/mL Blasticidin, and 1 µM Drug Selectable Marker 1 (DSM1) (Ghorbal et al., 2014). After parasites grew back up from transfection and were cloned using limiting dilution, clones were maintained in medium containing 50 nM aTc and 2.5 µg/mL Blasticidin. *Pf*J2^apt^-PDI11^glms^ parasites were generated in the same manner, using 50 µg pV5-glmS-PDI11 and 50 µg pUF1-Cas9-PDI11.

To generate the *Pf*PDI8^wt^ overexpression parasite line, RBCs were transfected with 50 µg pUC57-Hsp110-PDI8^wt^ and 50 µg pUF1-Cas9-Hsp110, then fed to *Pf*J2^apt^ parasites. Drug pressure was applied 48 hours after transfection, selecting with 0.5 µM aTc, 2.5 µg/mL Blasticidin, and 1 µM Drug Selectable Marker 1 (DSM1) (Ghorbal et al., 2014). After parasites grew back up from transfection and were cloned using limiting dilution, clones were maintained in medium containing 10 nM aTc and 2.5 µg/mL Blasticidin. *Pf*PDI11^wt^ and *Pf*PDI11^mut^ parasites were generated in the same manner, using pUC57-Hsp110-PDI11^wt^ and pUC57-Hsp110-PD11^mut^, respectively.

Parasite synchronization was carried out as described (Kudyba et al., 2019).

### Western Blotting

Western blots were performed as previously described (Muralidharan et al., 2011). Briefly, ice-cold 0.04% saponin in 1x PBS was used to isolate parasites from host cells. Parasite pellets were subsequently solubilized in protein loading dye to which Beta-mercaptoethanol had been added (LI-COR Biosciences) and used for SDS-PAGE. Primary antibodies used in this study were rat-anti-HA 3F10 (Roche, 1:3000), mouse-anti-HA 6E2 (Cell Signaling Technology, 1:1000), rabbit-anti-HA 715500 (Thermofisher, 1:100), mouse-anti-V5 TCM5 (eBioscence, 1:1000), rabbit-anti-V5 D3H8Q (Cell Signaling Technology, 1:1000), rabbit anti-*Pf*BiP MRA-1246 (BEI resources, 1:500), rabbit-anti-*Pf*EF1α (from D. Goldberg, 1:2000), and mouse-anti-*Pf*PMV (from D. Goldberg 1:400). Secondary antibodies used were IRDye 680CW goat-anti-rabbit IgG and IRDye 800CW goat-anti-mouse IgG (Li-COR Biosciences, 1:20,000). Membranes were imaged using the Odyssey Clx Li-COR infrared imaging system (Li-COR Biosciences). Images of membranes were processed using ImageStudio, the Odyssey Clx Li-COR infrared imaging system software (Li-COR Biosciences). Densitometry analysis of western blot signal was also performed using ImageStudio (Li-COR Biosciences).

### Microscopy and Image Analysis

Parasites were fixed for IFA using 4% Paraformaldehyde and 0.03% glutaraldehyde, then permeabilized with 0.1% Triton-X100. Primary antibodies used were rat-anti-HA 3F10 (Roche, 1:100), mouse-anti-HA 6E2 (Cell Signaling Technology, 1:100), mouse-anti-V5 TCM5 (eBioscence, 1:100), rabbit-anti-V5 D3H8Q (Cell Signaling Technology, 1:100), and mouse-anti-*Pf*PMV (from D. Goldberg 1:1). Secondary antibodies used were Alexa Fluor 488 and Alexa Fluor 546 (Life Technologies, 1:100). Cells were mounted to slides using ProLong Diamond with DAPI (Invitrogen). Fixed and stained cells were imaged using a DeltaVision II microscope system with an Olympus IX-71 inverted microscope. Images were collected as a Z-stack and deconvolved using SoftWorx, then displayed as a maximum intensity projection. Images were processed using Adobe Photoshop, with adjustments made to brightness and contrast for display purposes.

For imaging of parasite cultures using light microscopy, aliquots of culture were smeared onto glass slides and field-stained using Hema3 Fixative and Solutions (Fisher Healthcare), which is comparable to Wright-Giemsa staining. Slides were imaged using a Nikon Eclipse E400 microscope with a Nikon DS-L1-5M imaging camera. To measure parasite size, images were taken and parasites measured using ImageJ (NIH).

### Growth Assays using Flow Cytometry

Aliquots of parasites culture were incubated in 8 µM Hoescht 33342 (Thermofisher Scientific) for 20 minutes at room temperature, then fluorescence was measured using a CytoFlex S (Beckman Coulter, Hialeah, Florida). Flow cytometry data were analyzed using FlowJo software(Treestar, Inc., Ashland, Oregon). For IC_50_ experiments, data were analyzed using the 4-parameter dose-response-curve function of Prism (GraphPad Software, Inc.).

### Immunoprecipitation Assays

Anti-HA immunoprecipitation (IP) assays were performed as previously described, using anti-HA magnetic beads (Pierce) (Fierro et al., 2020). Anti-V5 IP assays were performed in the same manner as with anti-HA, but anti-V5 magnetic beads were used according to manufacturer instructions (MBL International Corporation).

### Mass Spectrometry and Data Analysis

CoIP samples were sent to Emory University Integrated Proteomics Core and analyzed using a Fusion Orbitrap mass spectrometer, or to the proteomics core at the Fred Hutchinson Cancer Research Center, where samples were analyzed using an OrbiTrap Elite. Data were searched using Proteome Discoverer 2.2 with UP000001450 *Plasmodium falciparum* (Uniprot Nov 2018) as the background database. The validation also included Sequest HT and Percolator to search for common contaminants. Results consisted of high confidence data with a 1% false discovery rate. Protein abundance was calculated by summing the total intensities (MS1 values) of all matched peptides for each selected protein, and normalizing by the total summed intensity of all matched peptides in the sample, as previously described (Boucher et al., 2018).

### Identification of PfJ2 Redox Partners

*Pf*J2^apt^ parasites were incubated with 3 mM divinyl sulfone (DVSF, Fisher Scientific) in 1x PBS for 30 minutes at 37°C, then used for anti-HA immunoprecipitation as described above. Immunoprecipitated proteins were separated by SDS-PAGE. Polyacrylamide gel slices corresponding to the protein molecular weights of interest were excised and the peptides extracted by in-gel enzymatic digestion. The gel slices were dehydrated in 100% acetonitrile and dried using a speed vac. The proteins were then reduced by rehydrating the gel slices in 10mM dithiothreitol in 100mM ammonium bicarbonate solution, and alkylated in 50mM iodoacetamide in 100 mM ammonium bicarbonate solution. The gel slices were then washed in 50% acetonitrile in 50mM ammonium bicarbonate solution before being dehydrated and dried again using 100% acetonitrile and a speed vac. Proteins were then digested in-gel by rehydrating the gel slices in trypsin enzyme solution consisting of 6ng/µl trypsin (Promega) in 50mM ammonium bicarbonate solution. Digestion was performed at 37°C overnight. Peptides were extracted through stepwise incubations with 2% acetonitrile and 1% formic acid solution, 60% acetonitrile and 0.5% formic acid solution, and 100% acetonitrile solution. Supernatants were combined and dried in a speed vac before resuspension in 20 µl water with 0.1% formic acid.

LC-MS/MS analysis was performed using a 50 cm fused silica capillary (75 µm ID) packed with C18 (2 µm, Dr. Maisch GmbH), and heated to 50°C. Prior to loading the column, sample was loaded onto a 2 cm Acclaim PepMap 100 (Thermo Fisher Scientific) trap (75 µm ID, C18 3 µm). For each sample injection, 5 µl of sample was loaded onto the trap using an Easy nLC-1000 (Thermo Fisher Scientific). Each sample was separated using the Easy nLC-1000 with a binary mobile phase gradient to elute the peptides. Mobile phase A consisted of 0.1% formic acid in water, and mobile phase B consisted of 0.1% formic acid in acetonitrile. The gradient program consisted of three steps at a flow rate of 0.3 µL/min: (1) a linear gradient from 5% to 40% mobile phase B over two hours, (2) a 10 minute column wash at 80% mobile phase B, and (3) column re-equilibration for 20 minutes at 5% mobile phase B.

Mass spectra were acquired on a Fusion Lumos Tribrid (Thermo Fisher Scientific) mass spectrometer operated by data dependent acquisition (DDA) using a top 15 selection count. Precursor ion scans were performed at 120,000 resolution over a range from 375 to 1375 m/z. DDA was performed with charge exclusion of 1 and greater than 8, with isotope exclusion, and dynamic exclusion set to 10 seconds. MS/MS was performed using an isolation window of 1.6 m/z for selection, normalized collision energy (NCE) of 28, and higher energy collision induced dissociation (HCD). MS/MS spectra were acquired at 15000 resolution with an automatic gain control (AGC) target of 70,000 and maximum injection time of 50 ms.

Mass spectra (.raw files) were converted to mzML format using MSConvert (version 3.0.1908) (Adusumilli & Mallick, 2017) and peptide sequences were identified using database searching with Comet (Eng et al., 2013) (version 2016.01 rev 2). Spectra were searched against a subset of *P. falciparum* secretory proteins, common laboratory contaminants, and an equal number of randomized decoy sequences (4386 total protein sequence). Comet parameters included variable modifications of +57.021464 Da or +118.0089 Da on cysteine and a variable modification of +15.994915 Da on methionine or tryptophan. Precursor mass tolerance was set to 25 ppm and a fragment_bin_tolerance of 0.02 and fragment_bin_offset of 0 were used. Full-tryptic enzymatic cleavage was set, allowing for up to 3 missed cleavages. Peptide spectrum matches (PSM) were analyzed using the Trans-Proteomic Pipeline (Deutsch et al., 2015) (TPP, version 5.0.0 Typhoon), to assign peptide and protein probabilities using PeptideProphet (Keller et al., 2002) and iProphet (Shteynberg et al., 2011), respectively. Spectral counts and precursor ion intensities were exported for each non-redundant PSM at a 1% false discovery rate (FDR). Protein inference was performed with ProteinProphet (Nesvizhskii et al., 2003), using a 1% FDR. The mass spectrometry proteomics data have been deposited to the ProteomeXchange Consortium via the PRIDE partner repository (Vizcaino et al., n.d.) with dataset identifier PXD019100.

## Acknowledgements

We thank Dan Goldberg for antibodies against *Pf*PMV and *Pf*EF1α; Anat Florentin for comments on the manuscript; Julie Nelson at the CTEGD Cytometry Shared Resource Laboratory for help with flow cytometry and analysis; and Muthugapatti Kandasamy at the Biomedical Microscopy Core at the University of Georgia for help with microscopy. We acknowledge assistance of the Proteomics Resource at Fred Hutchinson Cancer Research Center and the Emory University Integrated Proteomics Core for mass spectrometry and data analysis. This work was supported by awards from the ARCS Foundation to D.W.C., and the US National Institutes of Health (R01AI130139) to V.M. and (T32AI060546) to D.W.C. We are grateful for support from the National Institutes of Health from the Office of The Director, under award number S10OD026936, and by National Institute of General Medical Sciences under award numbers R01GM087221 and P41GM103533.

## Competing Interests

The authors declare no competing interests.

## Competing Interests

The authors declare no competing interests.

## Supplementary Figures

**Supplementary Figure 1.**
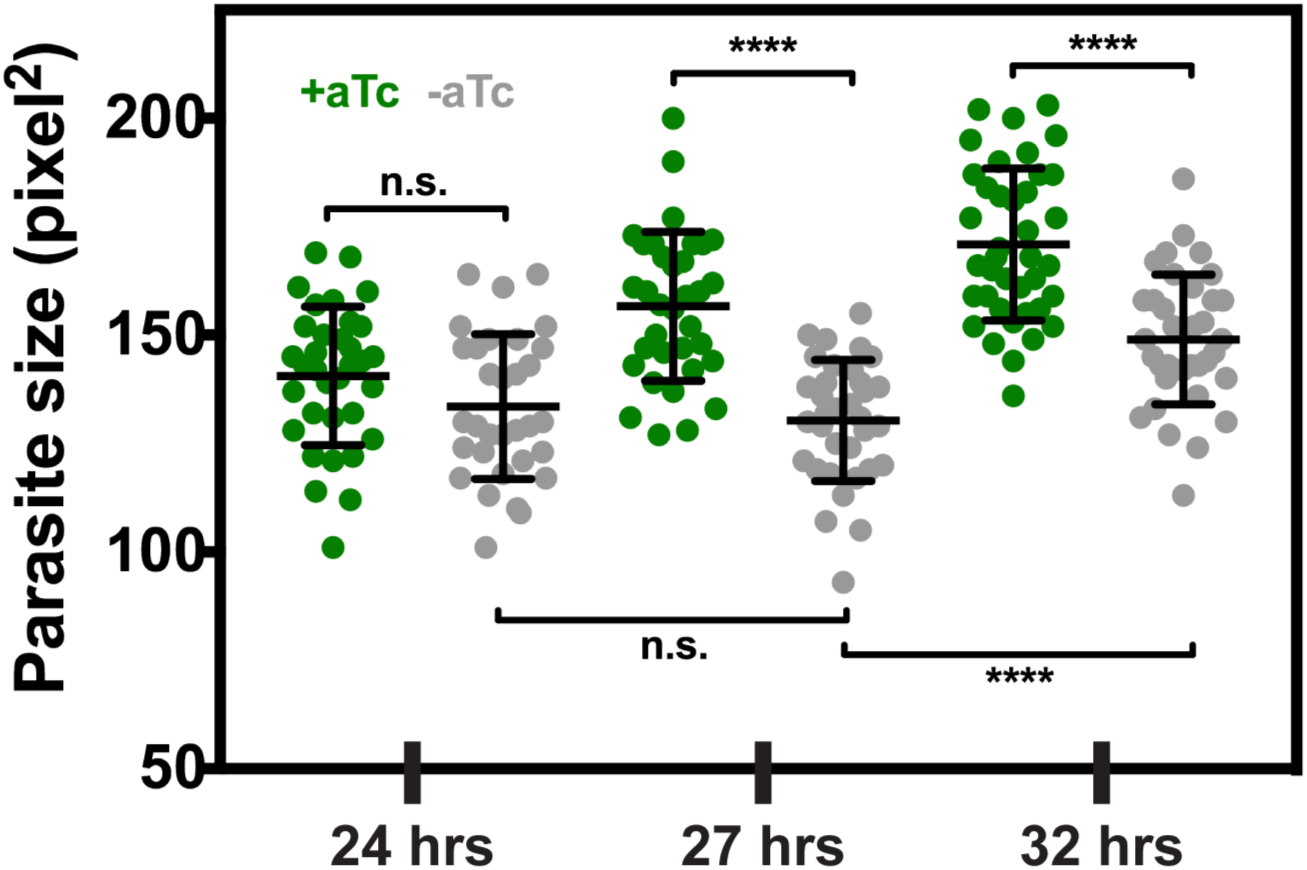
Parasite development is slowed during PfJ2 knockdown. *Pf*J2^apt^ parasites were tightly synchronized (0-3 hours) to the ring stage, then split into either +aTc (10 nM) or –aTc medium. Smears were made and field-stained at various time points throughout the asexual lifecycle. Stained slides were imaged and parasite size was measured. Unpaired t-test, **** indicates p ≤ 0.0001. Representative experiment of 3 biological replicates shown.

**Supplementary Figure 2.**
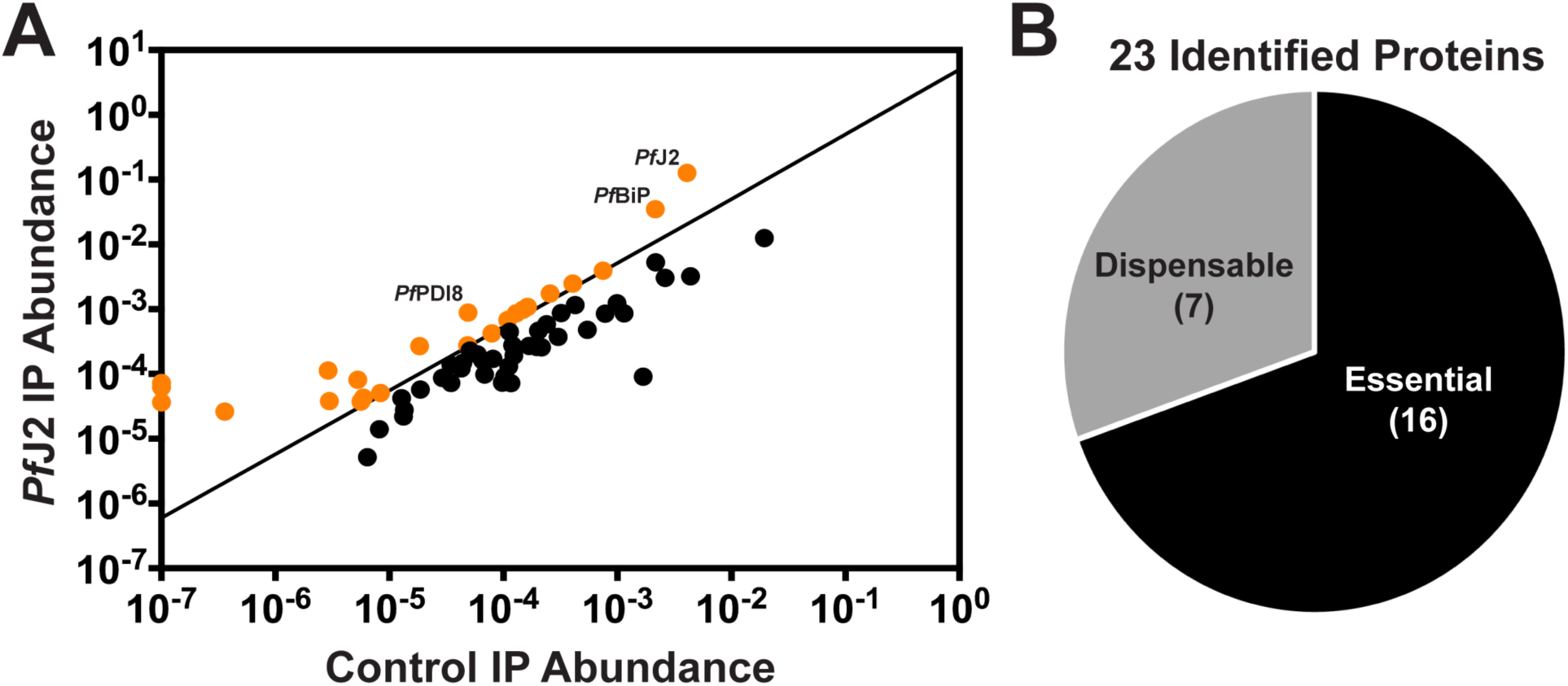
*Pf*J2 interacts with other essential chaperones, proteins in the secretory pathway. **A)** *Pf*J2 was immunoprecipitated from *Pf*J2^apt^ parasites using anti-HA antibody, and co-immunoprecipitated proteins were identified by tandem mass spectrometry analysis. Control, parental parasites were also used for immunoprecipitation and analyzed in the same manner. Each coIP experiment was performed in triplicate, and the abundance of each identified protein was calculated as previously described in Boucher *et al*. Candidate proteins of interest were further identified as those in the secretory pathway (predicted to contain a signal peptide and/or transmembrane domains) and those which were present in all three *Pf*J2^apt^ replicates and demonstrated a 5-fold enrichment compared to control experiments (shown in orange). **B)** The 23 proteins meeting our strict criteria were assessed against the piggyBac mutagenesis screen performed by Zhang, Wang *et al*., and 16 were predicted to have essential functions in the *P. falciparum* asexual stages.

**Supplementary Figure 3.**
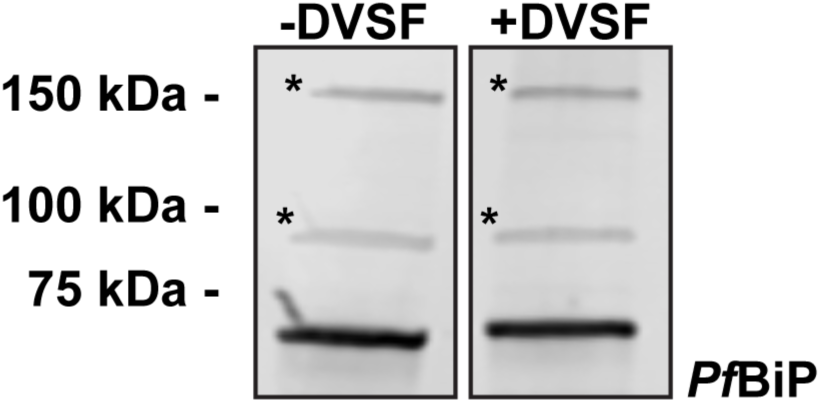
PfBiP SDS-PAGE migration is unaffected by DVSF. *Pf*J2^apt^-PDI8^glms^ parasites were treated with 3 mM DVSF in 1xPBS for 30 minutes at 37°C, or left untreated as a control, and parasite lysates were used for western blotting. Membranes were probed with antibodies against *Pf*BiP. Asterisks (*) denote nonspecific bands.

**Supplementary Figure 4.**
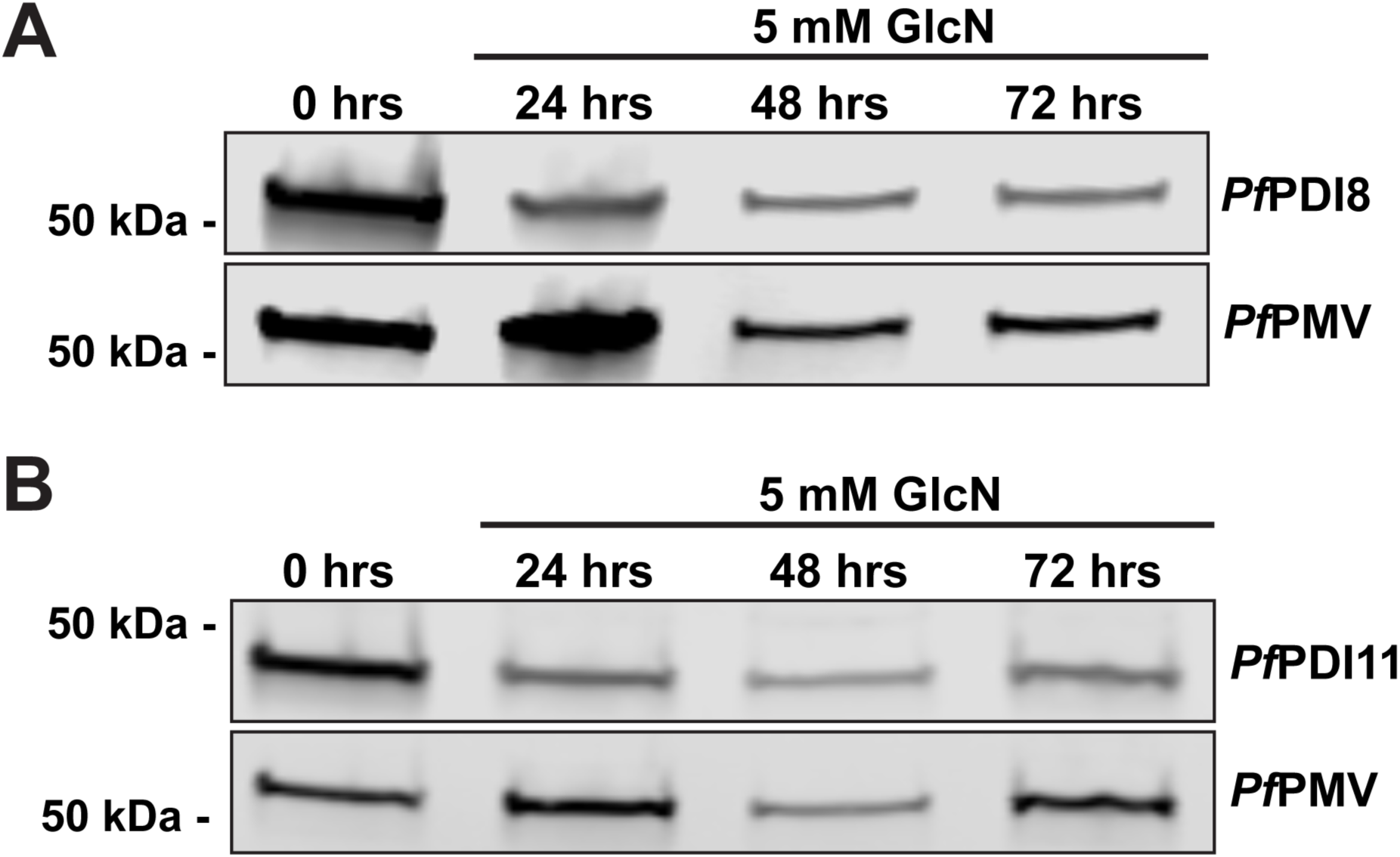
GlcN treatment leads to knockdown of *Pf*PDI8 and *Pf*PDI11. Asynchronous *Pf*J2^apt^-PDI8^glms^ (top) and *Pf*J2^apt^-PDI11^glms^ (bottom) parasites were treated with 5 mM GlcN and samples were taken for western blot analysis at 0 (before addition of GlcN), 24, 48, and 72 hours. Membranes were probed with antibodies for V5 and *Pf*PMV.

**Supplementary Figure 5.**
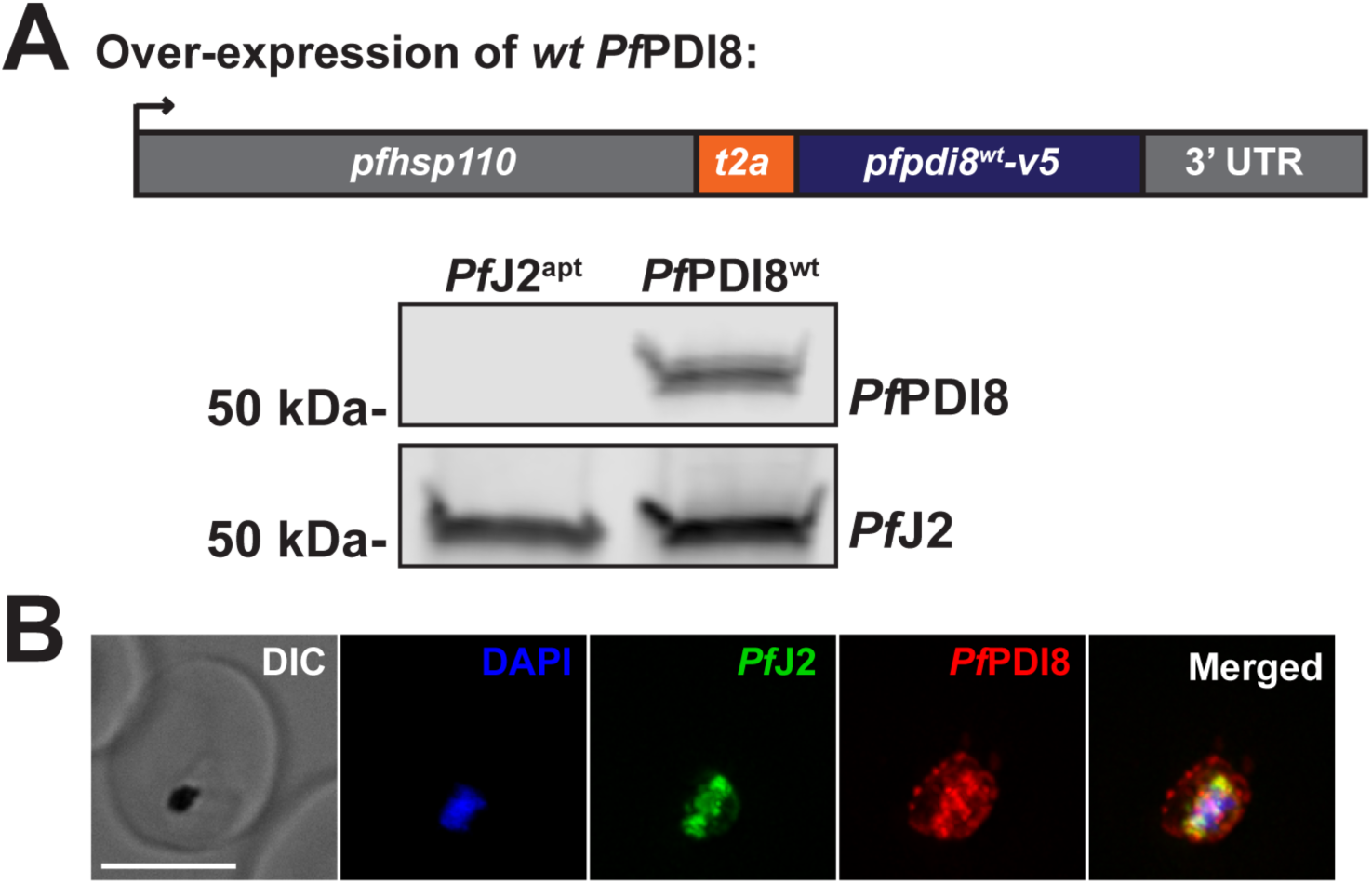
Overexpression of *Pf*PDI8 results in mislocalization. **A)** Top: schematic of exogenous V5-tagged, wild-type *Pf*PDI8 expression using the *pfhsp110* (PF3D7_ 0708800) locus and a T2A skip peptide in the *Pf*PDI8^wt^ parasite line, created in the background of *Pf*J2^apt^ parasites. Bottom: western blot of parental *Pf*J2^apt^ and *Pf*PDI8^wt^ parasite lysates, probed for antibodies against the V5 tag (*Pf*PDI8) and the HA tag (*Pf*J2). **B)** *Pf*PDI8^wt^ parasites were glutaraldehyde/paraformaldehyde fixed and used for IFA. Staining was carried out using DAPI (blue), antibodies against the HA tag (*Pf*J2, green), and the V5 tag (*Pf*PDI8, red). Scale bar represents 5 μm.

**Supplementary Figure 6.**
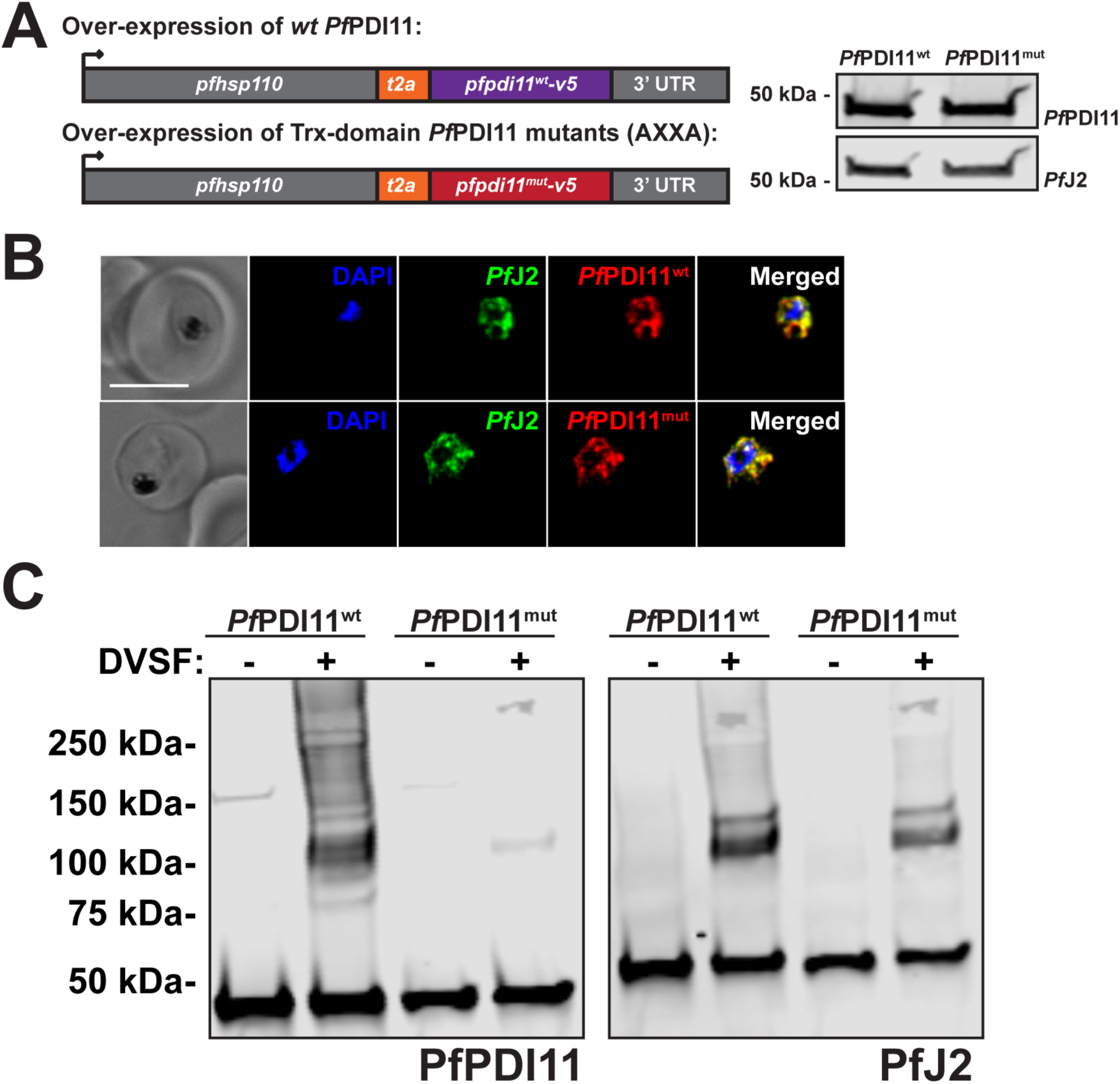
Characterization of PfPDI11 overexpression lines. **A)** Left: schematic of exogenous V5-tagged, *Pf*PDI11 expression using the *pfhsp110* (PF3D7_ 0708800) locus and a T2A skip peptide in the *Pf*PDI11^wt^ and *Pf*PDI11^mut^ parasite lines, created in the background of *Pf*J2^apt^ parasites. In *Pf*PDI11^mut^ parasites, both Trx-domain CXXS active sites were changed to AXXA. Right: western blot of *Pf*PDI11^wt^ and *Pf*PDI11^mut^ parasite lysates, probed for antibodies against the V5 tag (*Pf*PDI11) and the HA tag (*Pf*J2). **B)** *Pf*PDI11^wt^ and *Pf*PDI11^mut^ parasites were glutaraldehyde/paraformaldehyde fixed and used for IFA. Staining was carried out using DAPI (blue), antibodies against the HA tag (*Pf*J2, green), and the V5 tag (*Pf*PDI8, red). Scale bar represents 5 μm**. C)** *Pf*PDI11^wt^ and *Pf*PDI11^mut^ parasites were treated with 3 mM DVSF in 1x PBS for 30 minutes at 37°C, then parasite lysates used for western blotting. Membranes were probed with antibodies against the V5 tag (*Pf*PDI11) and the HA tag (*Pf*J2).

**Supplementary Figure 7.**
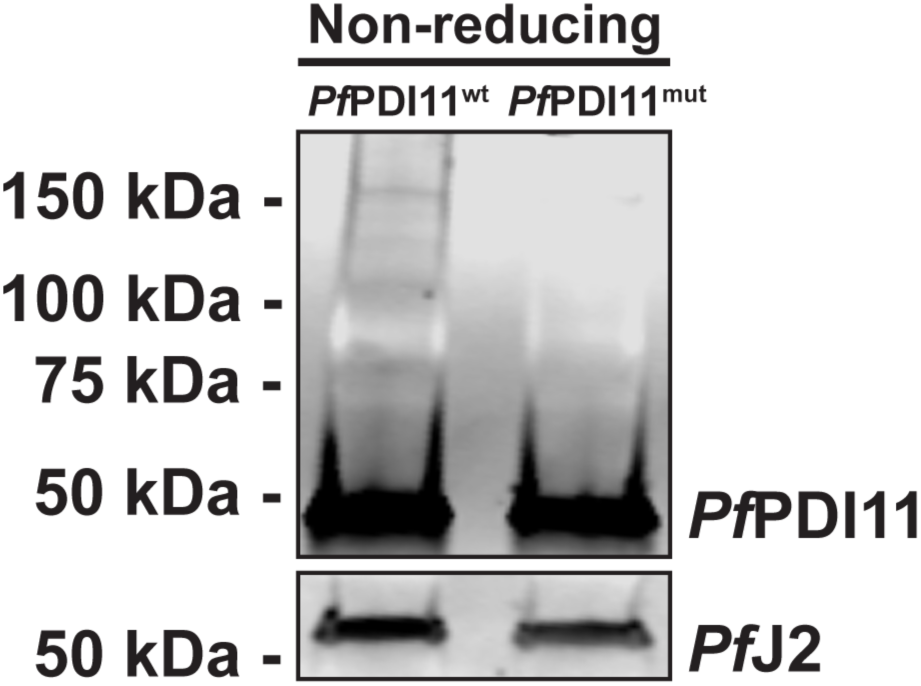
PfPDI11 non-reducing western blots. Saponin-isolated *Pf*PDI11^wt^ and *Pf*PDI11^mut^ parasites were dissolved in protein loading dye lacking a reducing agent and used for western blotting. Membranes were probed with antibodies against the V5 tag (*Pf*PDI11) and the HA tag (*Pf*J2).

**Supplementary Figure 8.**
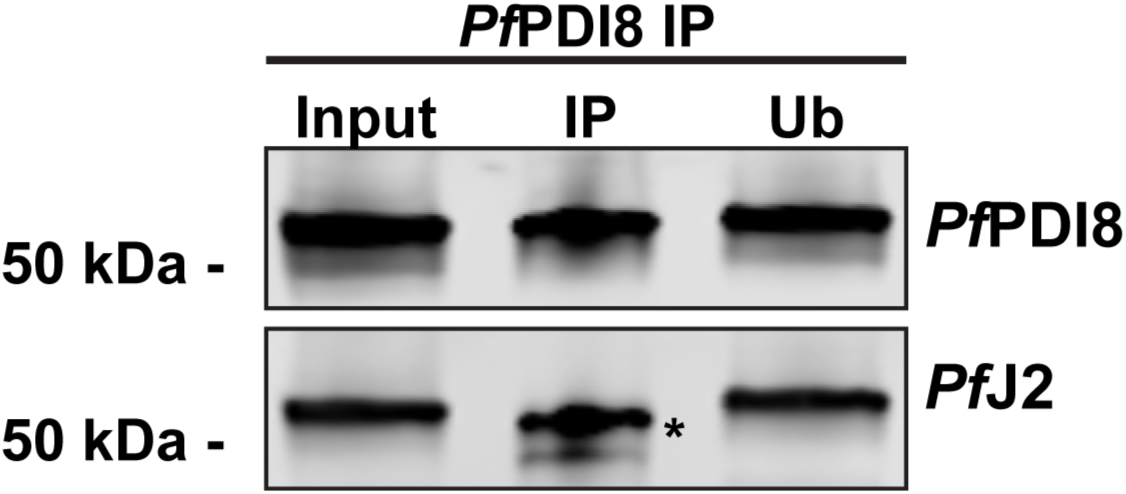
Immunoprecipitation of PfPDI8. *Pf*PDI8 and interacting proteins were immunoprecipitated from *Pf*J2^apt^-PDI8^glmS^ parasite lysate using anti-V5 antibodies. The pre-IP input sample, the sample eluted from the antibodies (IP), and the sample removed from the beads containing the unbound proteins (Ub) were used for western blotting. Membranes were probed with antibodies against V5 and HA. The asterisk (*) denotes the heavy chain of the antibody used for immunoprecipitation.

**Supplementary Figure 9.**
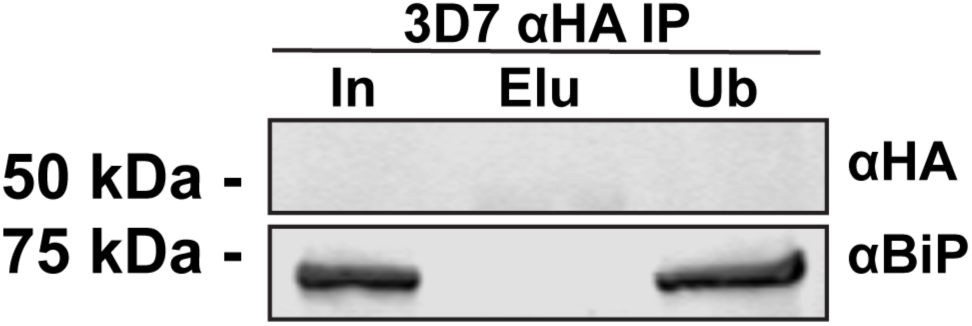
Anti-HA BiP IP. 3D7 parasite lysates were used for anti-HA immunoprecipitation (IP). The pre-IP input sample (In), the sample eluted from the antibodies (Elu), and the sample removed from the beads containing the unbound proteins (Ub) were used for western blotting. Membranes were probed with antibodies against HA and *Pf*BiP.

**Supplementary Figure 10.**
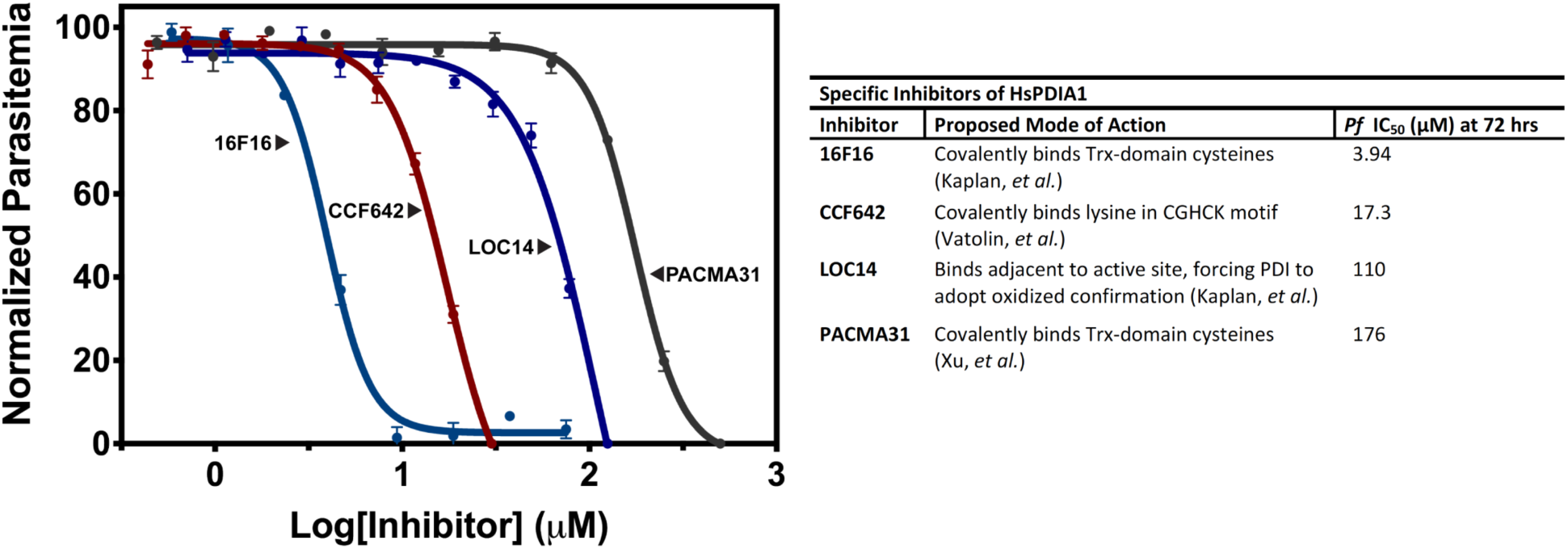
PDI Inhibitor IC50s. Left: Asynchronous 3D7 parasites were incubated in various concentrations of human PDI inhibitors. Each data point represents the mean parasitemia at a given concentration, in technical triplicate, at 72 hours. Error bars, which are not shown for data points in which they are smaller than the circle symbol, represent standard deviation from the mean. Representative IC_50_ curves are shown for each drug. Experiments were performed in biological triplicate. Right: table describing each of the human PDI inhibitors used and their calculated *P. falciparum* IC_50_ values.

## Notes

### Competing Interest Statement

The authors have declared no competing interest.

